# Control of *Clostridiodes difficile* virulence and physiology by the flagellin homeostasis checkpoint FliC-FliW-CsrA in the absence of motility

**DOI:** 10.1101/2022.11.08.515568

**Authors:** Duolong Zhu, Katherine J. Wozniak, Firas Midani, Shaohui Wang, Xingmin Sun, Robert A. Britton

## Abstract

Mutations affecting *Clostridioides difficile* flagellin (FliC) have been shown to be hypervirulent in animal models and display increased toxin production and alterations in central metabolism. The regulation of flagellin levels in bacteria is governed by a tripartite regulatory network involving *fliC*, *fliW*, and *csrA*, which creates a feedback system to regulate flagella production. Through genomic analysis of *C. difficile* clade 5 strains (non-motile), we identified they have jettisoned many of the genes required for flagellum biosynthesis yet retain the major flagellin gene *fliC* and regulatory gene *fliW*. We therefore investigated the roles of *fliC*, *fliW*, and *csrA* in the clade 5 ribotype 078 strain *C. difficile* 1015, which lacks flagella and is non-motile. Analysis of mutations in *fliC*, *fliW*, and *csrA* (and all combinations) on *C. difficile* pathogenesis indicated that FliW plays a central role in *C. difficile* virulence as animals infected with strains carrying a deletion of *fliW* showed decreased survival and increased disease severity. These *in vivo* findings were supported by *in vitro* studies showing that mutations impacting the activity of FliW showed increased toxin production. We further identified that FliW can interact with the toxin positive regulator TcdR, indicating that modulation of toxin production via FliW occurs by sequestering TcdR from activating toxin transcription. Furthermore, disruption of the *fliC*-*fliW*-*csrA* network results in significant changes in carbon source utilization and sporulation. This work highlights that key proteins involved in flagellar biosynthesis retain their regulatory roles in *C. difficile* pathogenesis and physiology independent of their functions in motility.

**IMPORTANCE:** *C. difficile* is a leading cause of nosocomial antibiotic-associated diarrhea in developed countries with many known virulence factors. In several pathogens, motility and virulence are intimately linked by regulatory networks that allow coordination of these processes in pathogenesis and physiology. Regulation of *C. difficle* toxin production by FliC has been demonstrated *in vitro* and *in vivo* and has been proposed to link motility and virulence. Here we show that clinically important, non-motile *C. difficile* strains have conserved FliC and regulatory partners FliW and CsrA, despite lacking the rest of the machinery to produce functional flagella. Our work highlights a novel role for flagellin outside of its role in motility and FliW in the pathogenesis and physiology of *C. difficile*.

## INTRODUCTION

*Clostridioides difficile* is a Gram-positive, spore-forming, toxin-producing, anaerobic bacterium that is a leading cause of nosocomial antibiotic-associated diarrhea in developed countries (1-3). Recently, *C. difficile* infection (CDI) reports from the community are rising and becoming a larger portion of total CDI cases (4). CDI can result in a spectrum of symptoms, ranging from mild diarrhea to pseudomembranous colitis and death (5). *C. difficile* possesses many virulence factors, such as toxins, binary toxins, adhesions, and flagella, among which toxin A (TcdA) and toxin B (TcdB) are the major ones (6, 7).

While flagella impact bacterial motility, biofilm formation, and host colonization, mutations in flagellin have been shown to play roles in virulence, toxin production, and fitness in several bacteria (8-10). The flagellin protein FliC has been reported to play important roles during *C. difficile* infection in motile strains (9, 11-14). Aubry et al. (11) reported that disruption of flagellar structural gene *fliC* resulted in increased toxin expression. Conversely, disruption of early-stage flagellar genes, such as *fliF*, *fliG*, or *fliM*, led to a significant reduction of toxin production in *C. difficile* 630Δ*erm*. Similarly, Baban et al. (12) reported that *tcdA* expression increased 44.4-fold in a *fliC* gene mutant, while *flgE* mutation led to a 10-fold reduction of *tcdA* expression in *C. difficile* 630Δ*erm*. Interestingly, deletion of other genes involved in flagellar assembly can abolish *C. difficile* motility completely but do not result in a significant change in toxin production that is observed in a *fliC* mutant (11). Based on previous studies, Stevenson et al. hypothesized that the regulation of the flagellar structural genes on toxin expression could be modulated by the direct change or loss of flagellar genes (such as *fliC* gene deletion) rather than the loss of the functional flagella and bacterial motility (9). Although the increase of pathogenicity of *fliC* mutant *in vivo* has been confirmed and corroborated by several studies (12, 13, 15), how *fliC* modulates *C. difficile* toxin production and pathogenesis remains unclear.

The partner-switching mechanism Hag-FliW-CsrA, which governs flagellin homeostasis and serves as a checkpoint of flagellar morphogenesis, was first described in *Bacillus subtilis* (16). In this system, Hag (flagellin protein and FliC homolog), FliW (a flagella synthesis regulator), and CsrA (a carbon storage regulator) form a sensitive regulatory feedback loop to regulate flagellum biosynthesis in *B. subtilis*. During this feedback regulation, FliW is first released from a FliW-Hag complex after the flagellar assembly checkpoint of hook completion and then binds to CsrA by a noncompetitive mechanism to relieve CsrA-mediated *hag* translation repression to increase flagellin synthesis concurrent with filament assembly (16, 17). CsrA is an ancestral protein and has evolved to modulate various physiological processes by regulating mRNA stability and/or translation initiation of target mRNA (18-24). CsrA typically binds to multiple specific sites that are located near or overlapping the cognate Shine-Dalgarno (SD) sequence in target transcripts (25, 26). In *B. subtilis*, two CsrA binding sites were identified in the 5’-UTR of *hag* transcripts (26). Oshiro et al. (27) further quantitated the interactions of FliW, Hag, and CsrA, and determined that Hag-FliW-CsrA^dimer^ functions at nearly 1:1:1 stoichiometry in *B. subtilis*.

The regulatory effects of *csrA* on bacterial physiology and carbon metabolism in *C. difficile* 630Δ*erm* were elucidated through the overexpression of the *csrA* gene (28). However, specific roles of FliW beyond its function as a flagellin synthesis regulator were not reported. We previously characterized the pleiotropic roles of FliW and CsrA in *C. difficile* R20291 (29), supporting a similar partner-switching mechanism termed FliC-FliW-CsrA. Nevertheless, more detailed investigations into the FliC-FliW-CsrA molecular mechanism and the modulation of the partner-switching mechanism on pathogenesis in *C. difficile* are needed.

Previous studies have shown that ribotype 078 (Clade 5) *C. difficile* strains lack flagella and bacterial motility (9). Surprisingly, FliC and FliW are highly conserved in the non-motile clade 5 strains of *C. difficile*, despite their lack of flagella and other genes encoding regulatory and structural genes involved in motility. Here, we investigated the roles of *fliC*, *fliW*, and *csrA* in modulating the pathogenesis and physiology of *C. difficile*, independent of flagellum biosynthesis. Our findings reveal that FliW plays a central role in *C. difficile* toxin production and virulence.

## RESULTS

### *fliC*, *fliW*, and *csrA* are conserved and expressed in non-motile *C. difficile* strains

We previously have shown that *fliW* and *csrA* genes are broadly found in *C. difficile* genomes and FliW and CsrA are highly conserved in divergent *C. difficile* ribotypes (29). In *C. difficile* R20291 (RT027 and motile), FliW and CsrA regulate flagellin (FliC) synthesis and bacterial virulence. We found that *fliC*, *fliW*, and *csrA* are conserved in non-motile *C. difficile* strains, although many other genes involved in flagellar synthesis have been lost (29). To evaluate the roles of *fliC*, *fliW*, and *csrA* in non-motile clade 5 *C. difficile* strains, we compared the flagellar regulon genes of RT078 (CD1015), RT033 (CLO_DA8024AA_AS), RT045 (CLO_EA6022AA_AS), RT126 (CLO_EA6160AA_AS), and RT127 (CLO_BA6185AA_AS) to the clade 2 RT027 (R20291). Clade 5 *C. difficile* strains completely lack early-stage flagellar genes (F3) and only the late-stage flagellar genes (F1, including *fliC*, *fliW*, and *csrA*) and glycosylation genes (F2) were found in their genomes (Fig. 1A, Fig. S1A, and Table S1). We selected RT078 strain CD1015 for further study and confirmed that CD1015 was non-motile (Fig. 1B). To confirm that FliC was still expressed in CD1015 despite the absence of flagella, we detected FliC in the cytoplasm of CD1015 and motile strain R20291 (Fig. 1C). We verified the cotranscription of *fliW* and *csrA* (Fig. S1B) and measured the transcription of *fliC* and *fliW*-*csrA* (Fig. S1C). These results showed that the *fliC*, *fliW*, and *csrA* genes are expressed in non-motile *C. difficile*, prompting us to explore the roles of these three genes in *C. difficile* pathogenesis, independent of their roles in flagellar biosynthesis and motility.

**Fig. 1.**
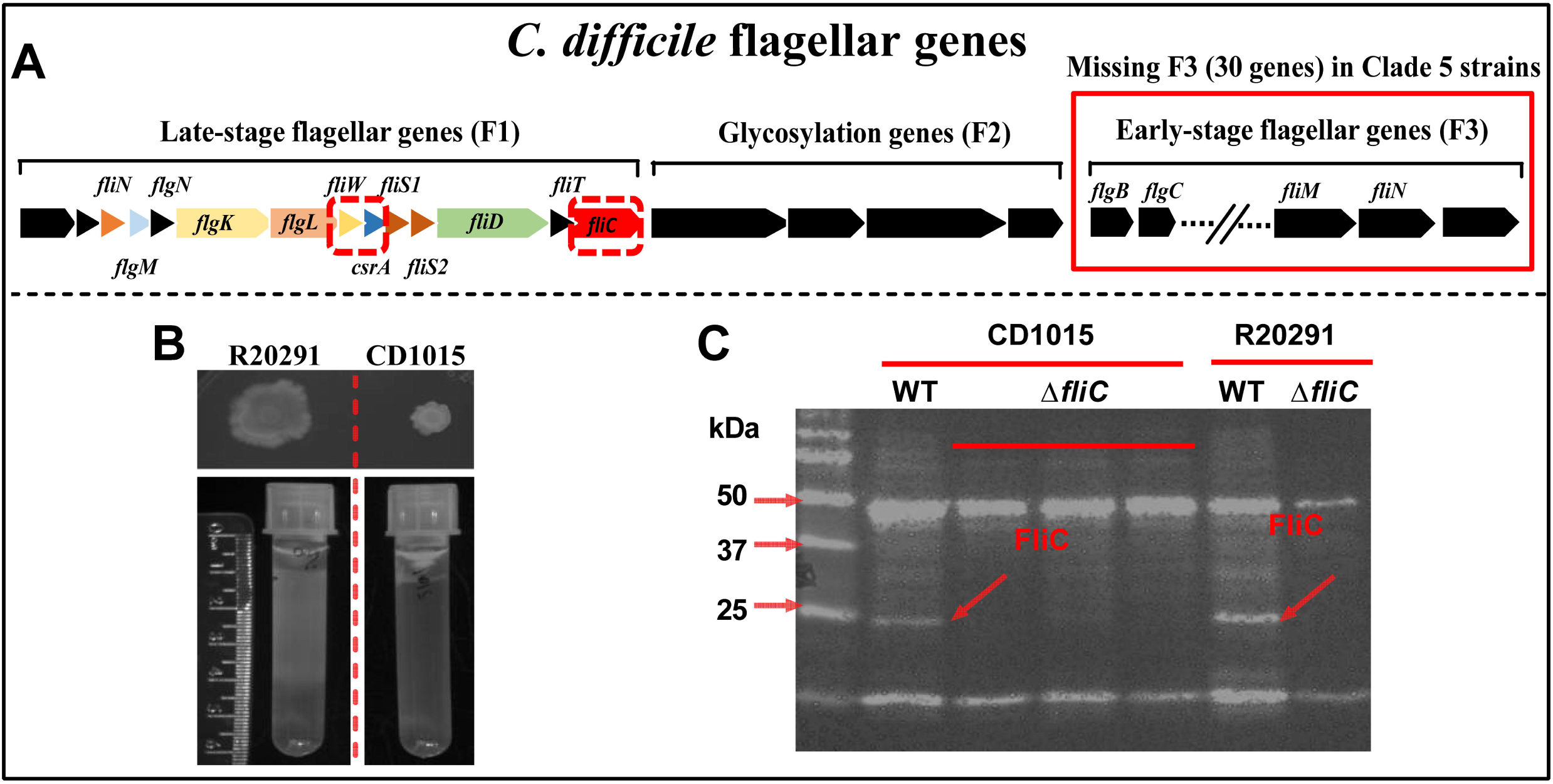
Non-motile clade 5 *C. difficile* conserves and expresses *fliC*, *fliW*, and *csrA*. **(A)** Schematic representation of conserved late-stage flagellar (F1) and glycosylation (F2) genes in the non-motile clade 5 ribotype 078 *C. difficile* strain 1015. **(B)** Comparison of CD1015 and R20291 motility. **(C)** Detection of FliC with anti-FliCD antibody in CD1015 and R20291 by Western blotting.

### FliC-FliW-CsrA network negatively regulates *C. difficile* toxins production

To test whether *fliC*, *fliW* and/or *csrA* modulate toxin production and bacterial pathogenicity in non-motile *C. difficile*, we constructed single deletions of *fliC*, *fliW*, *csrA* as well as all double mutant combinations and the triple mutant in CD1015 using a CRISPR-Cas12a system (Fig. S2A) (30, 31). Growth profiles of the different mutants were assessed in BHIS and TY media, and no significant differences in growth nor motility were observed (Fig. S2B-D). Supernatants of *C. difficile* cultures grown in TY media at 72 h post-inoculation were collected and toxin A and B levels were measured by ELISA. All mutants displayed significantly increased toxin production and transcription, with the notable exception of the Δ*fliC*Δ*csrA* double mutant, which produced wild-type levels of toxins A and B (Fig. 2A-B).

**Fig. 2.**
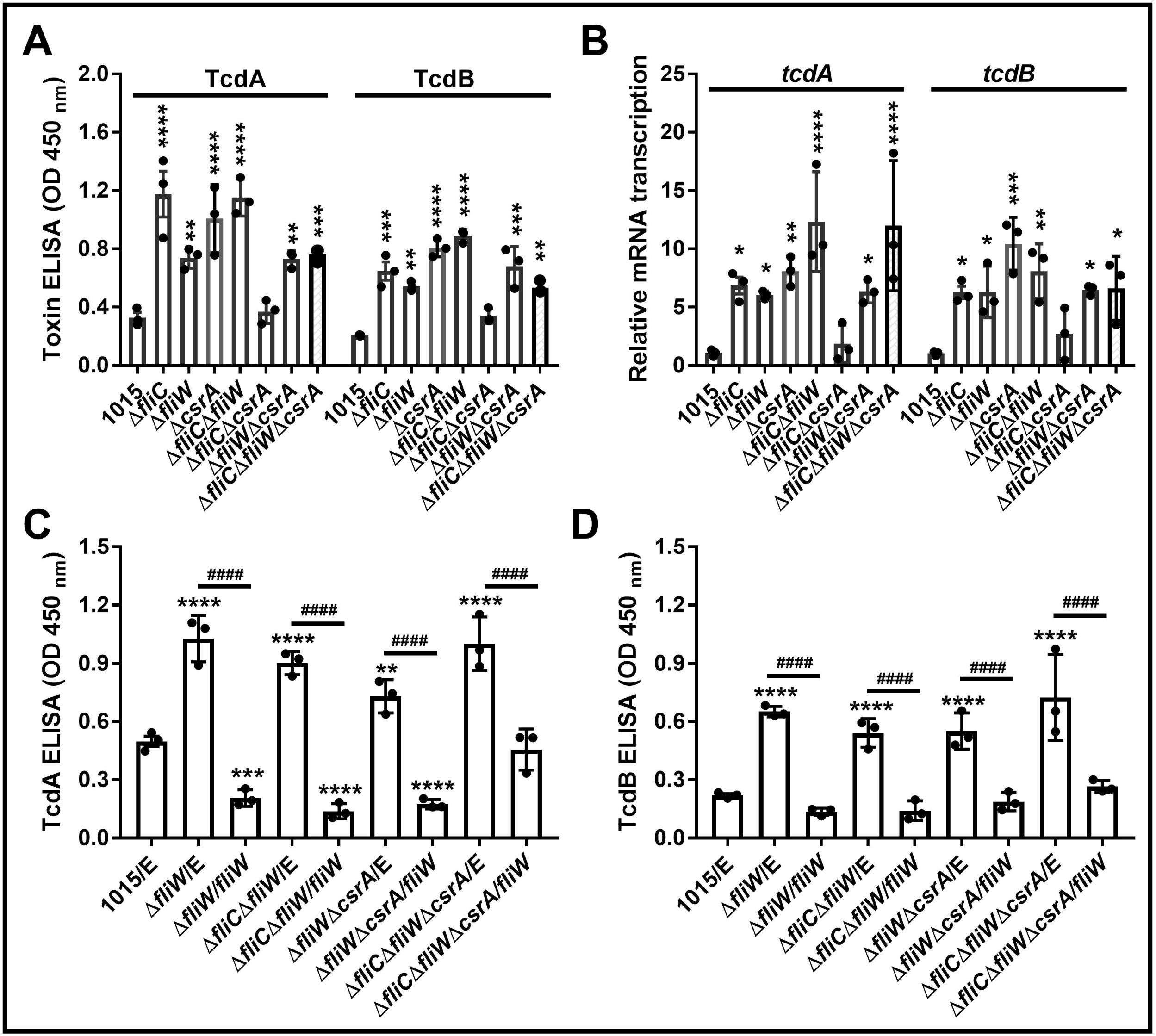
FliC-FliW-CsrA network negatively regulates *C. difficile* toxins production. **(A)** Toxin concentration in the supernatants of CD1015 and its derivative mutants detected by ELISA. **(B)** Transcription of *tcdA* and *tcdB* in CD1015 and its derivative mutants detected by RT-qPCR. **(C)** TcdA production in *fliW* complementation mutants. **(D)** TcdB production in *fliW* complementation mutants. The supernatants of different cultures were collected at 48 h of post inoculation in TY media, following analyzed by Toxin ELISA kits at OD_450 nm_. Bars stand for mean ± SEM. * means the significant difference of experimental strain compared to CD1015 (A-B) or CD1015 with empty plasmid (C-D) (\**P* < 0.05, \*\**P* < 0.01, \*\*\**P* < 0.001, \*\*\*\**P* < 0.0001). ^#^ means the significant difference of *fliW* complementation strain compared to mutant with empty plasmid (C-D) (^####^*P* < 0.0001). Differences were considered statistically significant if *P* < 0.05. One-way ANOVA with post-hoc Tukey test was used for statistical significance.

Interestingly, all mutants carrying a deletion of *fliW* gene displayed increased toxin production. To determine whether *fliW* was sufficient to restore toxin concentrations to normal levels in several mutants, we provided *fliW* in trans to Δ*fliW*, Δ*fliC*Δ*fliW*, Δ*fliW*Δ*csrA*, and Δ*fliC*Δ*fliW*Δ*csrA* strains and measured toxin production. Our data indicate that *fliW* complementation effectively reduced toxin production either to wild-type levels or below (Fig. 2C-D). We note that overexpressing *fliW* in wild-type CD1015 can also reduce toxin A and B levels (Fig. S3), further supporting a role for FliW in negative regulation of toxin production. These data indicate that FliW is a key node in negatively regulating *C. difficile* toxin production.

### FliW is a key modulator of *C. difficile* pathogenesis in a non-motile strain

To evaluate the role of the FliC-FliW-CsrA system on CD1015 virulence *in vivo*, we utilized the mutant strains described above in a mouse model of CDI and measured their impact on disease severity. Antibiotic treated mice were orally challenged with CD1015, Δ*fliC*, Δ*fliW*, Δ*csrA*, Δ*fliC*Δ*fliW*, Δ*fliC*Δ*csrA*, Δ*fliW*Δ*csrA*, or Δ*fliC*Δ*fliW*Δ*csrA* vegetative cells (10^8^ CFU/mouse). As shown in Fig. 3A, mice infected with Δ*fliC*Δ*fliW* (log-rank analysis, *P* 0.001), Δ*fliW* (*P* 0.01), Δ*fliW*Δ*csrA* (*P* 0.01), or Δ*fliC*Δ*fliW*Δ*csrA* (*P* 0.05) showed a dramatic increase in mortality (70-90%) compared to CD1015. The single mutant strains, Δ*fliC* and Δ*csrA*, displayed an intermediate phenotype in which 30-40% of the mice succumbed to infection. The Δ*fliC*Δ*csrA* double mutant displayed no increase in disease severity, consistent with the lack of increase in toxin production of this strain *in vitro*. We scored the severity of disease daily using a previously described clinical sickness scoring system (CSS) based on behavioral changes, stool characteristics, and weight loss (32). The highest CSS score for each mouse within 7 days post-infection was used for CSS comparison between the CD1015 and mutant strains. Our results showed that clinical scores of all derivative mutants, except Δ*fliC*Δ*csrA,* were higher than that of the parent strain, indicating more severe disease in mice colonized mutants that disrupt FliC-FliW-CsrA regulation (Fig. 3B). These data support FliW as a key player (negative regulator) in the regulation of toxin production in this regulatory cascade.

**Fig. 3.**
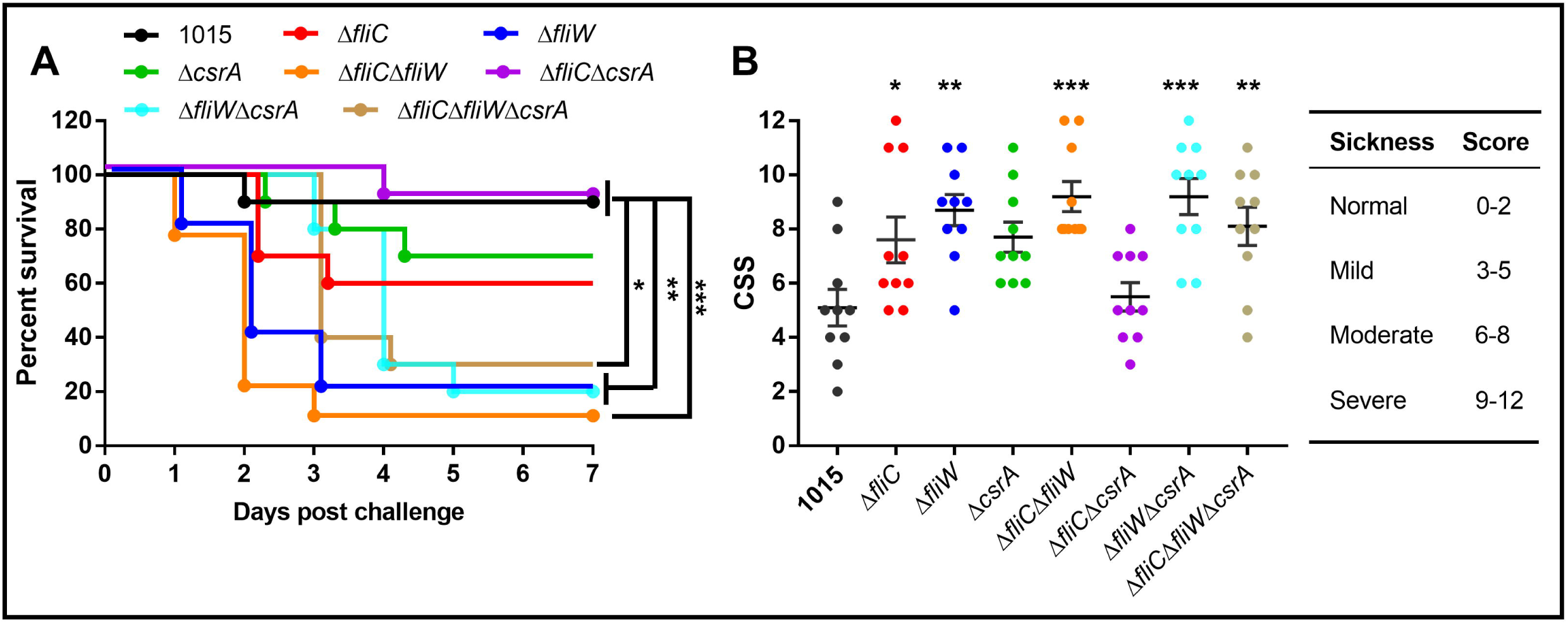
FliW is a key modulator of *C. difficile* pathogenesis in a non-motile strain. **(A)** Survival curve of CD1015 and its derivative mutants in a CDI mouse model. Animal survivals (10 mice/group) were analyzed by Kaplan-Meier survival analysis and compared by the Log-Rank test. Differences were considered statistically significant if *P* < 0.05 (\**P* < 0.05, \*\**P* < 0.01, \*\*\**P* < 0.001). **(B)** Disease severity in mice. The clinical sickness scoring system was used to evaluate the mice disease severity. The highest CSS score for each mouse within 7 days post infection was used for CSS comparison in different *C. difficile* strains. Animal experiments were repeated two times, data shown here were from a representative replicate. * means the significant difference of experimental strain compared to CD1015 (\**P* < 0.05, \*\**P* < 0.01, \*\*\**P* < 0.001). One-way ANOVA with post-hoc Tukey test was used for statistical significance.

### Characterization of protein-protein interactions of FliC-FliW-CsrA feedback network

Building on the partner-switching mechanism described in *B. subtilis* for this regulatory network, we wanted to explore the interactions governing the partner-switching mechanism FliC-FliW-CsrA in *C. difficile* (29). The *B. subtilis* model posits that FliC directly interacts with FliW, FliW can bind and antagonize the function of CsrA (Fig. 4A). To test whether these interactions occur in *C. difficile*, we performed a series of experiments to explore these interactions.

**Fig. 4.**
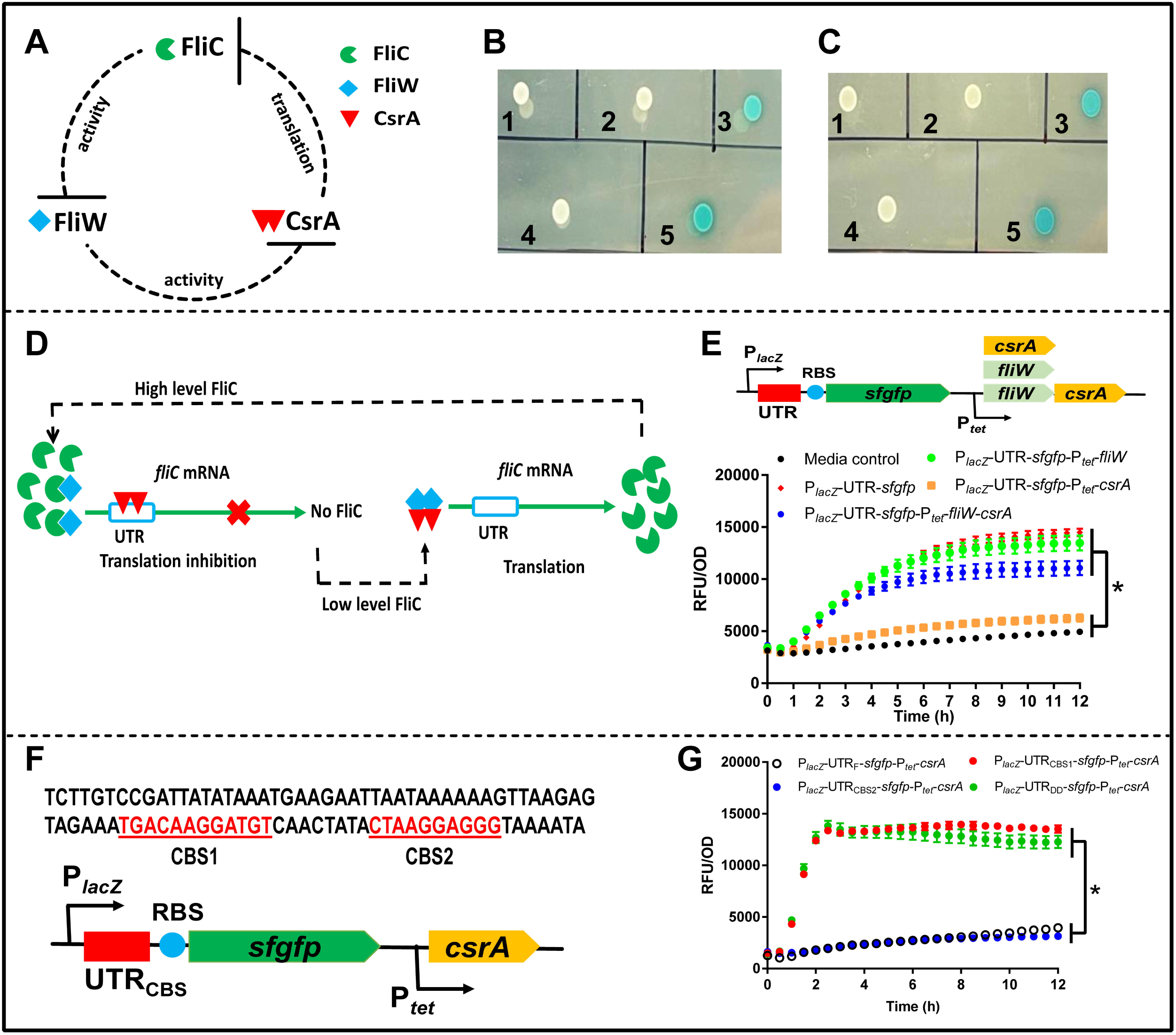
Characterization of FliC-FliW-CsrA regulatory loop in *C. difficile*. **(A)** Schematic representation of direct interactions of FliC, FliW, and CsrA in *C. difficile*. In the proposed FliC-FliW-CsrA regulatory loop, FliC binds FliW and FliW binds CsrA. **(B)** Protein-protein interaction test of FliC-FliW. 1: FliW-T25+T18; 2: FliC-T18+T25; 3: FliW-T25+FliC-T18; 4: T25+T18; 5: T25-Zip+T18-Zip. **(C)** Protein-protein interaction test of FliW-CsrA. 1: FliW-T25+T18; 2: CsrA-T18+T25; 3: FliW-T25+CsrA-T18; 4: T25+T18; 5: T25-Zip+T18-Zip. 2 µl of each reporter strain were spotted onto LB plates supplemented with IPTG and X-gal for indication. **(D)** Schematic representation of CsrA post transcriptional regulation on *fliC* transcripts. **(E)** Verification of post transcriptional regulation of CsrA on 5’UTR of *fliC* transcripts. Reporter plasmid pET21b-P*_lacZ_*-UTR-*sfgfp*-P*_tet_*-*csrA* (or *fliW* or *fliW*-*csrA*) was constructed, following the fluorescence of reporter strains with different recombinant plasmids were determined. **(F)** Prediction of potential <u>C</u>srA <u>b</u>inding <u>s</u>ites (CBS) in 5’-UTR of *fliC*. CBS1 and CBS2: CsrA potential binding sites 1 and 2 predicated by RNAstructure dynalign results analysis. **(G)** Identification of required binding sites for CsrA post-transcriptional regulation in 5’UTR of *fliC*. Three truncated 5’UTRs of *fliC* were assembled into the reporter plasmid, respectively, and subsequently the fluorescence of reporter strains with different truncated 5’UTR plasmids were analyzed. Statistical significance was determined at *P* < 0.05 (*), using a paired t-test.

We initially examined the interactions between the proteins FliW and FliC, as well as FliW and CsrA, using the *E. coli* two-hybrid (BACTH) system (33). We created C-terminal fusions of T25 fragment to FliW (FliW-T25) and T18 fragment to FliC (FliC-T18) and CsrA (CsrA-T18), respectively. As shown in Fig. 4B and 4C, we detected direct interactions between FliW and FliC (FliW-T25+FliC-T18) and between FliW and CsrA (FliW-T25+CsrA-T18) with no interaction detected in the negative controls, as predicted by the model and consistent with previous work in *B. subtilis*.

### Post-transcriptional regulation of *fliC* expression by CsrA

To investigate the post-transcriptional regulation of CsrA on *fliC* expression (Fig. 4D), we constructed a reporter plasmid (referred to as P*_lacZ_*-UTR-*sfgfp*-P*_tet_*-*csrA*) containing two regulatory elements: one is an IPTG-inducible promoter (P*_lacZ_*) driving the expression of the 5’ untranslated region of *fliC* (UTR) fused upstream of the reporter *sfgfp* gene (P*_lacZ_*-UTR-*sfgfp*); the other is a tetracycline-inducible promoter (P*_tet_*) driving the expression of *csrA* (P*_tet_*-*csrA*) (Fig. 4E). The strain containing only P*_lacZ_*-UTR-*sfgfp* was used as a positive control (P*_lacZ_*-UTR-*sfgfp*). As shown in Fig. 4E, induction of *csrA* along with UTR-*sfgfp* expression (P*_lacZ_*-UTR-*sfgfp*-P*_tet_*-*csrA*) decreased the fluorescence of the reporter strain dramatically compared to that of P*_lacZ_*-UTR-*sfgfp*, indicating negative regulation of CsrA on *fliC* expression. Meanwhile, *fliW*-*csrA* coexpression (P*_lacZ_*-UTR-*sfgfp*-P*_tet_*-*fliW*-*csrA*) could restore fluorescence signal to a level similar to the positive control strain, indicating FliW antagonizes CsrA. These results support a model in which CsrA negatively regulates *fliC* expression by interaction with the UTR of *fliC* and FliW antagonizes the negative regulation by CsrA. To exclude the impact of CsrA on IPTG inducible *fliC* transcription, we measured *sfgfp* transcription in different reporter strains containing either P*_lacZ_*-UTR-*sfgfp*, P*_lacZ_*-UTR-*sfgfp*-P*_tet_*-*fliW*, P*_lacZ_*-UTR-*sfgfp*-P*_tet_*-*csrA*, or P*_lacZ_*-UTR-*sfgfp*-P*_tet_*-*fliW*-*csrA* regulation cassette. No significant differences in *sfgfp* transcription were detected (Fig. S4A).

In *B. subtilis*, two CsrA binding sites have been identified in the 91 bp of the 5’ UTR of the *hag (fliC)* transcript (34). We compared *C. difficile fliC* and *B. subtilis hag* 5’-UTR RNA structures using the algorithm RNAstructure (35) and found two potential **<u>C</u>**srA **<u>b</u>**inding **<u>s</u>**ites (CBS1: from -45 to -34; CBS2: from -20 to -9, overlapping *fliC* Shine-Dalgarno sequence) included in two distinct predicted hairpin structures (Fig. 4F and Fig. S4B). To test whether the predicted binding sites are required for CsrA post-transcriptional regulation, we deleted either CBS1, CBS2, or both CBS1 and CBS2 (DD: **<u>d</u>**ouble binding sites **<u>d</u>**eletion) in the *fliC* 5’-UTR and assembled the full-length UTR (UTR_F_) and the truncated UTRs into the reporter plasmid (P*_lacZ_*-UTR_CBS_-*sfgfp*-P*_tet_*-*csrA*). As shown in Fig. 4G, CsrA lost its ability to regulate *fliC* post-transcriptionally when CBS2 (P*_lacZ_*-UTR_CBS1_-P*_tet_*-*csrA*) or CBS1-CBS2 (P*_lacZ_*-UTR_DD_-P*_tet_*-*csrA*) were deleted but kept similar regulation in the CBS1 deletion reporter strain (P*_lacZ_*-UTR_CBS2_-P*_tet_*-*csrA*). These results indicated that CBS2 (RBS) is required for CsrA post-transcriptional regulation on *fliC* expression. Taken together, we showed that *fliC* is expressed in non-motile *C. difficile*, characterized the partner-switching mechanism FliC-FliW-CsrA in *C. difficile*, and identified that CBS2 (RBS) is required for CsrA post-transcriptional on *fliC* expression.

### FliW interacts with toxin expression positive regulator TcdR

The alternative sigma factor TcdR is a key positive regulator of toxin production in *C. difficile*. As it not only positively regulates toxin expression but also affects *C. difficile* sporulation (36), this motivated us to investigate whether FliW interacts with TcdR. We first created C-terminal fusion of T18 fragment to TcdR (TcdR-T18) in the pUT18 plasmid and detected the direct interaction between FliW and TcdR using the BACTH system described above (Fig. 5A). To quantify the interaction strength between FliW and TcdR, we conducted a β-galactosidase assay using cell lysates from the reporter strain (BTH containing FliW-T25+TcdR-T18) by detecting the OD_420nm_ change over one hour after adding the substrate ONPG (ortho-Nitrophenyl-β-galactoside). As shown in Fig. 5B, although the control construct TcdR-T18+T25 showed a weak background color change, the FliW-T25+TcdR-T18 construct exhibited a 2-fold increase in the rate of color change compared to the TcdR-T18+T25 construct, indicating interaction between FliW and TcdR. Overall, our data suggest that FliW interacts with TcdR, which may antagonize TcdR’s positive role in the regulation of *C. difficile* toxin expression.

**Fig. 5.**
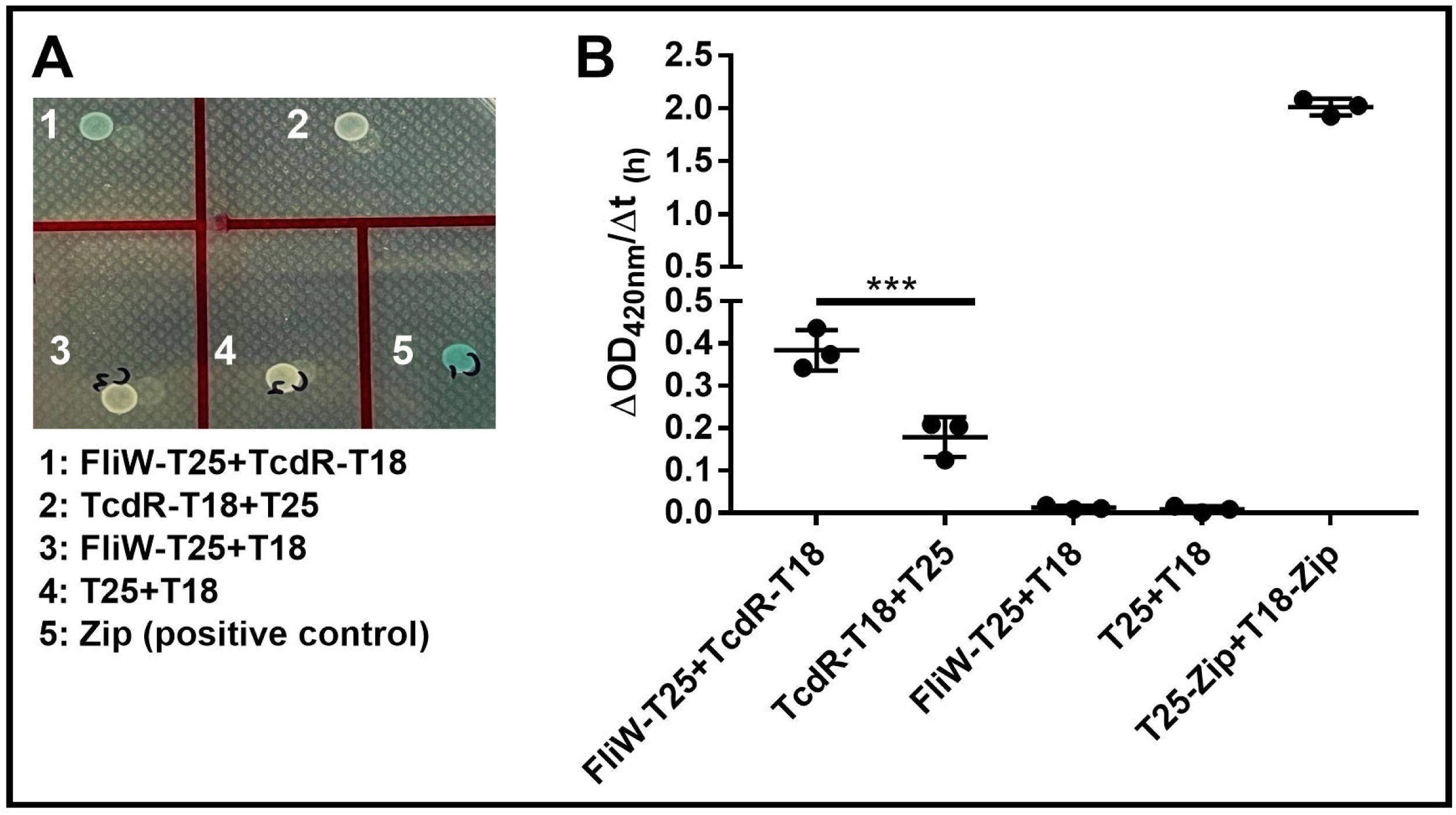
FliW interacts with the toxin expression positive regulator TcdR. **(A)** Protein-protein interaction test between FliW and TcdR *in vitro* using a bacterial two-hybrid system. **(B)** Determination of FliW and TcdR interaction strength. To quantify the interaction strength between FliW and TcdR, 10^10^ CFU of cell cultures were collected and lysed by bead beating. The cell lysates were used for β-galactosidase assay by detecting the OD_420nm_ change over one hour after adding the substrate ONPG. The interaction strength was calculated by ΔOD_420nm_ /Δt_(1h)_. Reporter BTH101 containing TcdR-T18+T25, FliW-T25+T18, or T25+T18 were used as negative controls. Reporter BTH101 containing T25-Zip+ T18-Zip was used as a positive control. 2 µl of reporter strain were dotted on LB+IPTG+X-gal indication plates when needed. Differences were considered statistically significant if *P* < 0.05 (\*\*\**P* < 0.001). One-way ANOVA with post-hoc Tukey test was used for statistical significance.

### Disruption of FliC-FliW-CsrA affects CD1015 strain carbon utilization

Given the central role of CsrA in bacterial carbon metabolism and that carbon utilization genes are dramatically upregulated in *C. difficile* Δ*fliC* mutants (37-39), we measured the growth profile of Δ*fliC*, Δ*fliW*, and Δ*csrA* mutants on 190 carbon sources using Biolog PM1 and PM2A plates. As shown in Fig. 6A, the three single mutants showed significant growth changes in 13 carbon substrates. Mannose and trehalose displayed the largest increase in growth, therefore we further examined *C. difficile* growth profiles in the different concentrations of these two sugars. CD1015, Δ*fliC*, Δ*fliW* and Δ*csrA* mutants were cultured in CDMM media supplemented with 40 mM mannose or 20 mM trehalose, respectively. Our results (Fig. 6B-C) showed that mannose poorly supports CD1015 growth in CDMM medium, but all of the mutants demonstrated robust growth on mannose. In trehalose supplemented CDMM media, mutants displayed similar growth rates to CD1015, but all mutants showed significantly delayed cell lysis (Fig. 6D).

**Fig. 6.**
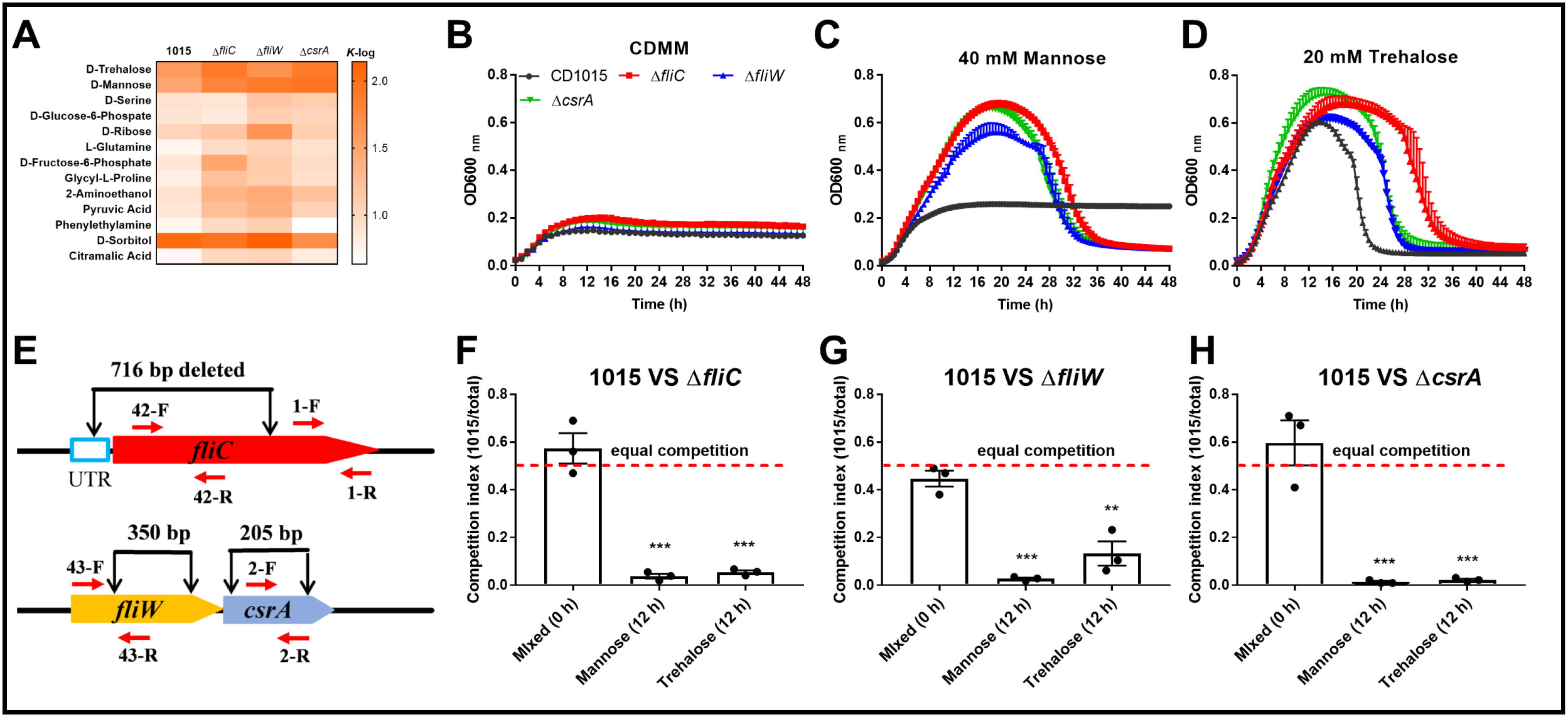
Disruption of FliC-FliW-CsrA affects CD1015 strain carbon utilization. **(A)** Carrying capacity [*K*, the maximum growth supported by the environment, *K*-log = ln(OD_max)-ln(OD_initial)] of growth profile of CD1015, Δ*fliC*, Δ*fliW*, and Δ*csrA* in Biolog PM1 and PM2A plates. **(B-D)** Growth curves of CD1015 and its derivative mutants in CDMM media without any added carbon source (B), 40 mM mannose (C), and 20 mM trehalose (D). **(E)** Primers used for the qPCR in the competition test. **(F-H)** Competition of CD1015 with Δ*fliC* (F), Δ*fliW* (G), and Δ*csrA* (H). Bars stand for mean ± SEM. * means the significant difference of CD1015 abundance in the competition mixtures compared to that of at 0 h (\*\**P* < 0.01, \*\*\**P* < 0.001). One-way ANOVA with post-hoc Tukey test was used for statistical significance.

To evaluate the impact of *fliC*, *fliW*, and *csrA* deletions on bacterial fitness in specific carbon sources, *in vitro* competition assays were performed. CD1015 and each mutant were co-cultured in CDMM media supplemented with either 40 mM mannose or 20 mM trehalose and incubated for 12 hours. Primers 1-F/R, which can anneal to both CD1015 and the mutants’ genomes, were used for total bacterial detection and primers 42-F/R, 43-F/R, and 2-F/R which can only pair to the wild-type genome were used for CD1015 detection (Fig. 6E). As shown in Fig. 6F-H, the abundance of CD1015 in competition with either Δ*fliC*, Δ*fliW*, or Δ*csrA* strains significantly decreased at 12 hours. The qPCR competition results were further verified by colony PCR test by isolating dozens of colonies from each competition (Fig. S5). Our competition data are consistent with the growth profile analysis, indicating that Δ*fliC*, Δ*fliW*, and Δ*csrA* mutants outcompete the wild type strain in trehalose and mannose supplemented media. This suggests that mutants might also outcompete the wild type *in vivo*, potentially impacting *C. difficile* pathogenesis.

## DISCUSSION

Bacteria possess complex regulatory networks that govern the decision to settle in a favorable environment or become motile to find new environments. In pathogens, the decision to make flagella or biofilms is often intimately tied to nutrient acquisition, virulence factor expression, and toxin production (40, 41). Clade 5 strains of *C. difficile*, which are phylogenetically distinct from the other four clades of *C. difficile* strains, are known to colonize livestock and cause human disease. Indeed, direct transmission between animals and humans has been described (42). Clade 5 strains are also unique in that they are non-motile and have lost most of the genes that encode for the machinery that builds and support the flagella, while retaining the major flagellin protein FliC (Fig. 1A and Fig. S1A). This prompted us to explore the role of the FliC-FliW-CsrA regulatory network on *C. difficile* physiology in the absence of motility and flagella formation.

The data presented here support that FliW is a key regulator that controls toxin production and virulence in CD1015 (RT078). Notably, mutation of *fliW* alone or in combination with *fliC* and/or *csrA* resulted in the greatest reduction in animal survival and increased disease severity score compared to CD1015 and other mutations. These findings correlated well with the amount of toxin A and B produced *in vitro*. We also found the double mutant Δ*fliC*Δ*csrA* was the only mutant that did not increase toxin production *in vitro* and did not result in increased disease severity *in vivo*. Finally, complementation data indicated that single *fliW* complementation can effectively reduce toxin production in Δ*fliW*, Δ*fliC*Δ*fliW*, Δ*fliW*Δ*csrA*, and Δ*fliC*Δ*fliW*Δ*csrA*, either to wild-type levels or below. These data support that FliW is the key node in negatively regulating toxin production and virulence in the FliC-FliW-CsrA regulatory network. Our previous work has also shown that *fliW* mutants cause increased toxin production in the motile hypervirulent *C. difficile* strain R20291 (29).

Previous studies have reported that *tcdR* mutants can affect both toxin production and sporulation in *C. difficile* (36), and that CsrA functions as a global post-transcriptional regulator in bacteria (19), involved in stage III sporulation in *Clostridium acetobutylicum* (43). Here, we not only verified that FliW binds to CsrA but also demonstrated that FliW can interact with TcdR, indicating that FliW can indirectly affect *C. dificile* toxin expression and sporulation (Fig. S6A). Because CsrA has been shown to regulate multiple aspects of central metabolism in many different bacterial species (19, 44), we also were interested in investigating whether the disruption of this regulatory network had an impact on carbon source metabolism. Recently, an *in vivo* gene expression study of R20291Δ*fliC* strain revealed that *treA*, encoding phosphotrehalase, increased 177-fold in the *fliC* mutant compared to the wild-type strain (13). Our previous study showed that dietary trehalose can contribute to the virulence of epidemic *C. difficile* (45). Strikingly, we observed that single mutations in all three genes (*fliC*, *fliW*, *csrA*) were able to confer the ability of CD1015 to grow on the sugar mannose and altered how trehalose is metabolized. *C. difficile* needs to balance sporulation, toxin production, and nutrient utilization in the infected host to successfully invade and colonize the host gut. In the future, one of the central questions moving forward with this line of research is to identify what additional inputs regulate the FliC-FliW-CsrA to impact *C. difficile* pathogenesis and physiology.

Elegant previous work has elucidated how the Hag-FliW-CsrA regulatory loop modulates flagellin homeostasis and bacterial motility in *B. subtilis* (26, 27). However, the export and assembly of flagella in *B. subtilis* is a key player in the complex partner switch mechanism of regulation, which is lost in CD1015 and other clade 5 *C. difficile* strains. Our data from two-hybrid studies and genetic experiments agree with the proposed model of direct interactions between FliC-FliW and CsrA-FliW with FliC being regulated post-transcriptionally by CsrA. Notably, in two predicted CsrA binding sites, only the Shine-Dalgarno sequence (located in CBS2) of *fliC* transcripts is required for CsrA post-transcriptional regulation, which is distinct from the regulation in *B. subtilis*.

Based on our findings presented in this paper, we hypothesize that deleting *csrA* could enhance FliC expression. This would increase FliC binding to FliW, reducing the amount of free FliW (no significant difference in *fliW* transcription between 1015 and Δ*csrA*) (Fig. S6B) in the cell. Consequently, more TcdR bound by FliW would be released, positively regulating toxin production. In the Δ*fliC* mutant, although no FliC is present to bind to FliW, we observed lower levels of *fliW* transcription (Fig. S6B). This is likely because no CsrA (*fliW*-*csrA* cotranscription) is needed for *fliC* translation repression, suggesting a potential reduction in FliW levels, which could also positively regulate toxin production. When both *fliC* and *csrA* were deleted, there was no disturbance observed in *fliW* transcription or the availability of FliW within the cell, resulting in no significant change in toxin production. We propose that FliW acts as a negative regulator of toxin production by antagonizing TcdR whose transcription or activity is regulated by FliC and CsrA. Disruption of the FliC-FliW-CsrA partner switch could result in less FliW available to participate in toxin repression (Fig. 7). Since FliW functions as a binding partner for several identified proteins, it would be worthwhile to uncover additional potential FliW binding targets using co-immunoprecipitation to better understand its roles in *C. difficile*.

**Fig. 7.**
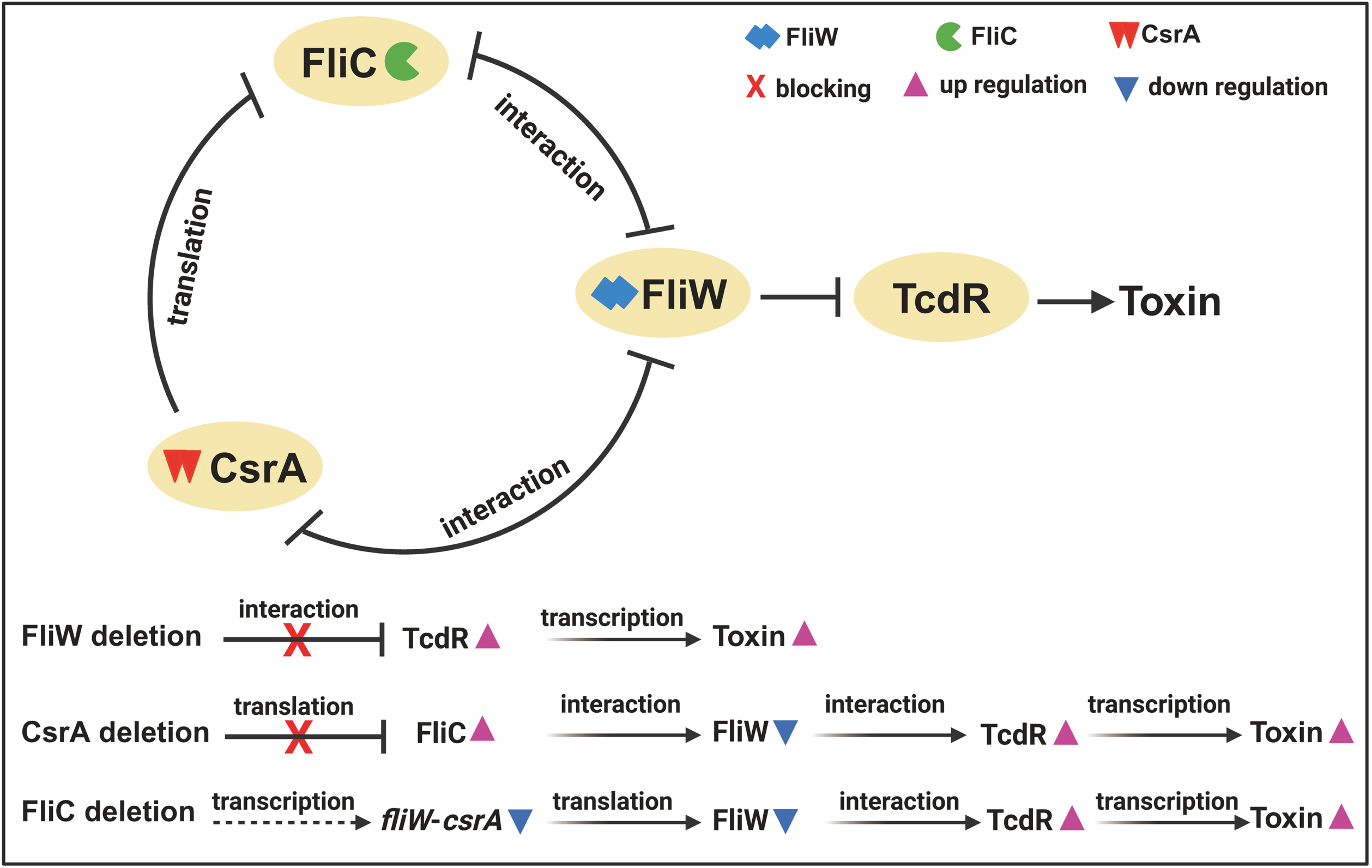
Regulation model of FliC-FliW-CsrA on CD1015 pathogenesis.

## MATERIALS AND METHODS

### Bacteria, plasmids, and culture conditions

Table S2 lists the strains and plasmids used in this study. *C. difficile* strains were cultured in BHIS media (brain heart infusion broth supplemented with 0.5% yeast extract and 0.1% L-cysteine, and 1.5% agar for agar plates) at 37 in an anaerobic chamber (90% N_2_, 5% H_2_, 5% CO_2_). For carbon substrate metabolization and Biolog analysis, *C. difficile* strains were cultured in CDMM media supplemented with specific carbon substrates (46, 47). For spores preparation,

*C. difficile* strains were plated on *C. difficile* 70:30 sporulation agar plates (6.3% Bacto peptone, 0.35% proteose peptone, 0.07% (NH_4_)_2_SO_4_, 0.106% Tris base, 1.11% BHI, 0.15% g yeast extract, 0.03% L-cysteine, 1.5% agar) and purified as described earlier (48). To enumerate *C. difficile* CFU in the fecal sample, fecal solutions were plated on adapted TCCFA plates which contain 5 μg/ml lysozyme and 50 μg/ml kanamycin (49). *Escherichia coli* DH5α, *E. coli* HB101/pRK24, *E. coli* BL21, and *E. coli* BTH101, were grown aerobically at 37 in LB media (1% tryptone, 0.5% yeast extract, 1% NaCl). *E. coli* DH5α was used as a cloning host, *E. coli* HB101/pRK24 was used as a conjugation donor host, *E. coli* BL21 was used as a protein expression host, and *E. coli* BTH101 was used as a protein-protein interaction analysis host. Antibiotics were added when needed: for *E. coli*, 20 μg/ml chloramphenicol, 50 μg/ml kanamycin, and 100 μg/ml ampicillin; for *C. difficile*, 15 μg/ml thiamphenicol, 250 μg/ml D-cycloserine, 50 μg/ml kanamycin, 16 μg/ml cefoxitin, and 500 ng/ml anhydrotetracycline.

### Chemicals and DNA manipulations

The DNA markers, PCR product purification kit, DNA gel extraction kit, restriction enzymes, cDNA synthesis kit, and SYBR Green RT-qPCR kit were purchased from Thermo Fisher Scientific (Waltham, USA). PCRs were performed with the high-fidelity DNA polymerase NEB Q5 Master Mix, and PCR products were assembled into target plasmids with NEBuilder HIFI DNA Assembly Master Mix (New England, UK). Primers (Table S3) were purchased from IDT (Coralville, USA). All chemicals were purchased from Sigma (St. Louis, USA) unless those stated otherwise.

DNA manipulations were carried out according to standard techniques (50). Gene *fliC*, *fliW*, and *csrA* were assembled into *EcoR*I-*BamH*I digested plasmid pMTL84153, resulting in pMTL84153-*fliC* (referred to as 153-*fliC*), pMTL84153-*fliW* (153-*fliW*), and pMTL84153-*csrA* (153-*csrA*) with primers 20-22, respectively. Gene edit plasmid pDL-1 containing Cas12a (AsCpfI) under control of tetracycline inducing promoter was used for *C. difficile* gene deletion according to the previous reports (30, 31). The target sgRNA was designed and analyzed on the Cas-OFFinder (http://www.rgenome.net/cas-offinder/) and CINDEL (http://big.hanyang.ac.kr/cindel/) websites. Plasmids were conjugated into *C. difficile* as described earlier (51). Briefly, constructed gene deletion plasmids (with primers 23-35) from *E. coli* DH5α were transformed into the donor host *E. coli* HB101/pRK24 and were conjugated into *C. difficile* strains subsequently. Transconjugants were selected on BHIS-TKC plates (15 μg/ml thiamphenicol, 50 μg/ml kanamycin, 16 μg/ml cefoxitin) and subcultured into BHIS-Tm broth (15 μg/ml thiamphenicol) to log phase, then the subsequent cultures were diluted with PBS serially and plated on induction plates (BHIS-Tm-ATc: 15 μg/ml thiamphenicol and 500 ng/ml anhydrotetracycline). After 48 h incubation, colony PCR was examined with check primers 1-C-F/R and 2-C-F/R to select the correct gene deletion colonies. Then the deletion plasmid was cured by several passages in BHIS broth without antibiotics. The genome of 1015Δ*fliC* (referred to as Δ*fliC*), 1015Δ*fliW* (Δ*fliW*), 1015Δ*csrA* (Δ*csrA*), 1015Δ*fliC*Δ*fliW* (Δ*fliC*Δ*fliW*), 1015Δ*fliC*Δ*csrA* (Δ*fliC*Δ*csrA*), 1015Δ*fliW*Δ*csrA* (Δ*fliW*Δ*csrA*), and 1015Δ*fliC*Δ*fliW*Δ*csrA* (Δ*fliC*Δ*fliW*Δ*csrA*) were isolated and used as templates for the PCR test with check primers, and the PCR products were sequenced to confirm the correct gene deletion.

### Growth profile and motility assay

*C. difficile* strains were cultured to an optical density of OD_600_ of 0.8 in BHIS or TY media and were diluted to an OD_600_ of 0.2. Then, 1% of the cultures were inoculated into fresh BHIS or TY, followed by measuring OD_600_ for 48 h.

To analyze the difference in carbon substrate utilization between parent strain and mutants, phenotype microarray plates (PM1 and PM2A) were used to conduct Biolog analysis. Briefly, *C. difficile* strains were cultured to an OD_600_ of 0.6 in BHIS, then 20% of cultures were inoculated into CDMM media with 0.5% glucose and were incubated for another 3 - 4 h (52). Following, 2 ml of cultures were centrifuged in the chamber and washed three times with CDMM base medium, and finally were resuspended in CDMM media and adjusted into the same OD for Biolog plates and specific carbon source (mannose and trehalose) utilization analysis. The Biolog plates containing bacteria were incubated in a plate reader at 37°C in an anaerobic chamber, with OD_600_ measurements taken every 10 minutes. The resulting data were automatically analyzed using AMiGA software developed by our lab (47).

To examine the effect of *fliC* deletion on CD1015 motility, CD1015 and CD1015Δ*fliC* were cultured to an OD_600_ of 0.8. For swarming analysis, 2 µl of cultures were dropped onto soft BHIS agar (0.2%) plates. For swimming analysis, 2 µl of *C. difficile* cultures were penetrated into a soft BHIS agar (0.15%) tube. The motility assay plates and tubes were incubated in the anaerobic chamber for 48 h.

### Toxin production, gene transcription, and bacteria competition assay

To evaluate toxin production in *C. difficile* strains, 10 ml of *C. difficile* cultures from different strains cultured in TY media were collected at 72 h post incubation. The cultures were adjusted to the same OD_600_ value, centrifuged at 4, 8000×*g* for 15 min, filtered with 0.22 μm filters, and then used for ELISA test (EAGLE, CDT35-K01).

For toxin transcription analysis (with primers 37-38), 2 ml of 48 h post inoculated *C. difficile* cultures in TY media were centrifuged at 4, 12000×*g* for 5 min for total RNA extraction. To assay *manA*, *treA*, and *treA2* gene expression (with primers 39-40), 5 ml of 8 h post inoculated *C. difficile* cultures in CDMM media supplemented with trehalose or mannose were used for RNA isolation with TRIzol reagent. The transcription of genes were measured by RT-qPCR. All RT-qPCRs were repeated in triplicate, independently. Data were analyzed by using the comparative CT (2^-ΔΔCT^) method with 16s rRNA as a control (with primers 36).

To compare the fitness of CD1015 and mutants in specific carbon substrates, *in vitro* competition assays were conducted. The same CFU of CD1015, Δ*fliC*, Δ*fliW*, and Δ*csrA* from the log phase were collected, centrifuged, and washed three times with CDMM base media in the anaerobic chamber. Following, 10^7^ CFU of CD1015 was mixed with the same CFU of Δ*fliC*, Δ*fliW*, and Δ*csrA*, respectively. The mixed seed cultures were then inoculated into CDMM media supplemented with mannose or trehalose. After 12 h of post-inoculation, 100 μl of mixed cultures were serially dilueted and plated on BHIS plates for colony PCR test with primers 3-C-F/R and 4-C-F/R. Meanwhile, the total genome from mixed cultures were isolated and detected by qPCR with primer pairs 1, 2, 42, and 43 for competition analysis. In the colony PCR test, CFU_1015_/CFU_mutant_ was calculated. In the qPCR test, the fold change of CD1015 accounting for total bacteria was calculated as: 2^(ΔCT^_12h_^-ΔCT^_0h_^)^, ΔCT= CT_1015_-CT_total_. Among them, CT_1015_ was acquired by primers 42-F/R in Δ*fliC*, or 43-F/R in Δ*fliW*, or 2-F/R in Δ*csrA* that only paired to the deleted gene sequence, and CT_total_ was acquired by primers 1-F/R paired to the remaining part of *fliC* gene after deletion. ΔCT_0_ was calculated with genome from the mixed seed cultures at 0 h.

### Bacterial virulence of mutants in the mouse model of CDI

C57BL/6 female and male mice (6 weeks old) were ordered from The Jackson Laboratory, Ellsworth, ME. All studies were approved by the Institutional Animal Care and Use Committee of Baylor College of Medicine. The experimental design and antibiotic administration were conducted as described earlier (53). Briefly, 80 mice were divided into 8 groups in 16 cages and each mouse was challenged with 10^8^ of *C. difficile* vegetative cells after orally administered antibiotic cocktail (kanamycin 0.4 mg/ml, gentamicin 0.035 mg/ml, colistin 0.042 mg/ml, metronidazole 0.215 mg/ml, and vancomycin 0.045 mg/ml) in drinking water for 6 days. After 6 days of antibiotic treatment, all mice were given autoclaved water for 2 days, followed by one dose of clindamycin (10 mg/kg, intraperitoneal route) 24 h before *C. difficile* challenge (Day 0). The mice were monitored daily for a week for changes in weight, diarrhea, stool characteristics, behavior, and mortality. All survived mice were humanely euthanized on day 7 of post *C. difficile* challenge.

Mice fecal pellets at post challenge days 1, 3, and 5 were collected from each mouse and were transferred into the anaerobic chamber as fast as possible. Following, the feces were diluted with PBS at a final concentration of 30 mg/ml and 100 μl of diluted feces were plated on TCCFA (5 μg/ml lysozyme and 50 μg/ml kanamycin added) plates to enumerate the total CFU of *C. difficile*. To calculate the spores CFU in the feces, the diluted feces solution were heated at 65 for 30 min, then plated on TCCFA plates as well. To evaluate toxin tilter in feces, 30 mg/ml of the fecal solution was used for TcdA and TcdB ELISA analysis.

### Protein-protein interaction analysis

To analyze the protein-protein interaction of FliC, FliW, and CsrA, a bacterial two-hybrid system (BACTH system kit) was used. We got the BACTH system kit as a gift from Dr. Bouveret (Aix-Marseille University, France) and followed the previously published protocol (33). Briefly, we digested plasmid pUT18 and pKNT25 with *BamH*I. Then *fliW*, *fliC*, *csrA,* and *tcdR* genes were amplified and assembled into the N-terminal of T25 and T18 tags with primers 5-10, respectively, resulting in plasmid pKNT25-*fliW* (referred as FliW-T25), pKNT25-*fliC* (FliC-T25), pKNT25-*csrA* (CsrA-T25), pUT18-*fliW* (FliW-T18), pUT18-*fliC* (FliC-T18), pUT18-*csrA* (CsrA-T18), and pUT18-*tcdR* (TcdR-T18). Following the two recombinant plasmids encoding FliW-T25 and FliC-T18 hybrid proteins were co-transformed into competent *E. coli* BTH101 reporter cells to analyze the interaction between FliW and FliC. For FliW and CsrA interaction analysis, reporter cells expressing FliW-T25 and CsrA-T18 hybrid proteins were used. Similarly, reporter cells containing FliW-T25 and TcdR-T18 were employed to test the interaction between FliW and TcdR. The correct cotransformants were first selected on LB plates with 100 μg/ml ampicillin and 50 μg/ml kanamycin. Then the protein-protein interaction was analyzed by dotting the recombinant BTH101 cultures on LB-X-Gal-IPTG indicator plates. The reporter strain containing empty plasmids pKNT25 and pUT18 was used as a control. The blue colonies indicate a positive result, while the white colonies indicate no interaction between two proteins.

To quantify the interaction strength between FliW and TcdR, we performed a β-galactosidase assay according to a previous report with several modifications (33). A total of 10^10^ CFU of cell cultures were collected and lysed by bead beating in 1 ml of Z buffer (8 g of Na_2_HPO_4_·12H_2_O, 3.125 g of NaH_2_PO_4_·H_2_O, 0.375 g KCl, 0.123 g MgSO_4_·7H_2_O dissolved in 500 ml H_2_O). The cell lysates were then centrifuged at 4, 12000×*g* for 10 min. Subsequently, 100 µl of the surenatants were used for the β-galactosidase assay by measuring the OD_420nm_ change after adding the substrate ONPG. The interaction strength was calculated as ΔOD_420nm_ /Δt_(1h)_. Reporter BTH101 containing TcdR-T18+T25, FliW-T25+T18, or T25+T18 were used as negative controls, while BTH101 containing T25-Zip+ T18-Zip was used as a positive control.

### Post-transcriptional regulation of CsrA on *fliC* expression

To investigate the post-transcriptional regulation of CsrA on *fliC* expression, we constructed several recombinant reporter plasmids that containing P*_lacZ_*-5’UTR-*sfgfp* [5’ untranslated region (5’-UTR) of *fliC* transcript and *sfgfp-*6-His (*sfgfp*) fluorescence reporter under the control of P*_lacZ_* induction promoter] and P*_tet_*-*csrA/fliW/fliW*-*csrA* (*csrA* or *fliW* or *fliW*-*csrA* under the control of tetracycline induction promoter P*_tet_*) regulation elements with primers 11-19 in pET21b plasmid using NEBuilder HIFI DNA Assembly Master Mix and transformed these reporter plasmids into the expression host *E. coli* BL21. To further identify the CsrA binding sites in 5’UTR of *fliC* transcripts, we analyzed and compared *fliC* and *hag* 5’-UTR structures on RNAstructure website (https://rna.urmc.rochester.edu/RNAstructureWeb/Servers/Predict2/Predict2.html), following we synthesized three truncated 5’UTR of *fliC* and assembled them into pET21b-P*_lacZ_*-5’UTR-*sfgfp*-P*_tet_*-*csrA* to evaluate the blocking efficiency of CsrA on *sfgfp* expression.

### Sporulation assay

*C. difficile* sporulation analysis were conducted as reported earlier (54). *C. difficile* strains were cultured on *C. difficile* 70:30 sporulation agar plates for 5 days, following bacterial lawns were scraped from plates and resuspended in PBS. Afterward, CFU of cultures resuspended in PBS was counted on BHIS plates with 0.1% TA for sporulation ratio analysis. The sporulation ratio was calculated as CFU (65 heated, 30 min) / CFU (not heated).

### Statistical analysis

The reported experiments were conducted in independent biological triplicates, and each sample was additionally taken in technical triplicates. Animal survivals were analyzed by Kaplan-Meier survival analysis and compared by the Log-Rank test. Student’s t-test was used for two groups comparison. One-way analysis of variance (ANOVA) with post-hoc Tukey test was used for more than two groups comparison. Results were expressed as mean ± standard error of the mean. Differences were considered statistically significant if *P* < 0.05 (*).

## ACKNOWLEDGMENTS

This work was supported by the National Institutes of Health (NIH) grants to R.A.B. (U01AI124290 and R01AI123278) and X.S (2R01AI132711). We thank Dr. Bouveret at Aix-Marseille University, France for gifting us the BACTH system kit.

## Supplementary figure legends

**Fig. S1 Conservation of flagellar genes in non-motile clade 5 strains and verification of *fliW*-*csrA* and *fliC* transcription.**

**(A)** Schematic representation of flagellar genes in the non-motile clade 5 strains and motile RT027 R20291. **(B)** Verification of *fliW* and *csrA* co-transcription by RT-PCR. M: DNA ladder; 1-3: CD1015 genomic DNA was used as PCR template; 4-6: CD1015 cDNA was used as PCR template. 1 and 4: *fliW* PCR test; 2 and 5: *csrA* PCR test; 3 and 6: *fliW*-*csrA* PCR test. **(C)** Transcription analysis of *fliC* and *fliW*-*csrA* in CD1015. Bars stand for mean ± SEM. Differences were considered statistically significant if *P* < 0.05 (\*\**P* < 0.01, \*\*\**P* < 0.001). One-way ANOVA with post-hoc Tukey test was used for statistical significance.

**Fig. S2 Generation of CD1015 derivative mutants, and test of bacterial growth profiles and motility.**

**(A)** Deletion of UTR-*fliC*, *fliW*, *csrA*, and *fliW*-*csrA*. 1-C-F/R were used to verify *fliC* deletion and 2-C-F/R were used to test *fliW*, *csrA*, and *fliW*-*csrA* deletion. M: DNA ladder; 1 and 3: CD1015 genome as PCR template; 2: CD1015Δ*fliC* (Δ*fliC*) genome test; 4: CD1015Δ*fliW* (Δ*fliW*) genome test; 5: CD1015Δ*csrA* (Δ*csrA*) genome test; 6: CD1015Δ*fliW*Δ*csrA* (Δ*fliW*Δ*csrA*) genome test. **(B)** Growth profile in BHIS media. **(C)** Growth profile in TY media. **(D)** Bacterial motility test. Swarming and swimming analysis were tested with soft BHIS agar (0.2%) plates and BHIS agar (0.15%) tubes, respectively.

**Fig. S3. Toxin expression in the *fliW* overexpression strain.**

TcdA and TcdB concentrations in the supernatants of CD1015-pMTL84153 (1015/E) and *fliW* overexpression strain (1015/*fliW*) were detected by ELISA. Bars stand for mean ± SEM. Differences were considered statistically significant if *P* < 0.05 (\**P* < 0.05). Statistical analysis was performed using an unpaired two-tailed *t* test.

**Fig. S4 Characterization of FliC-FliW-CsrA regulation loop in *C. difficile*.**

**(A)** Transcription of *sfgfp* in the different recombinant reporter strains. Red column: P*_lacZ_*-UTR-*sfgfp*; Green column: P*_lacZ_*-UTR-*sfgfp*-P*_tet_*-*fliW*; Orange column: P*_lacZ_*-UTR-*sfgfp*-P*_tet_*-*csrA*; Blue column: P*_lacZ_*-UTR-*sfgfp*-P*_tet_*-*fliW*-*csrA*; **(B)** Structure comparison of *B. subtilis* 5’-UTR of *hag* and CD1015 5’-UTR of *fliC*. BS1 and BS2: CsrA binding sites 1 and 2 in *B. subtilis*. CBS1 and CBS2: CsrA potential binding sites 1 and 2 in CD1015. RNAstructure dynalign results were calculated (calculate the lowest free energy secondary structures common to two unaligned sequences).

**Fig. S5 Δ*fliC*, Δ*fliW*, and Δ*csrA* mutants outcompete CD1015 in the presence of trehalose and mannose.**

**(A)** Primers used for the colony PCR in the competition test. **(B-D)** CFU_1015_/CFU_mutant_ in the competition cultures. * means the significant difference of CD1015 CFU in the competition mixtures compared to that of at 0 h (\**P* < 0.05, \*\**P* < 0.01). One-way ANOVA with post-hoc Tukey test was used for statistical significance.

**Fig. S6 Analysis of sporulation and *fliW* transcription in CD1015 and its derivative mutants**

**(A)** Sporulation analysis. *C. difficile* strains were cultured on 70:30 sporulation agar plates for 5 days, following which the scraped cultures were 10-fold diluted and plated on BHIS plates with 0.1% TA to detect sporulation ratio. The sporulation ratio was calculated as CFU (65 heated, 30 min) / CFU (not heated). **(B)** Comparison of *fliW* transcription in CD1015, Δ*fliC*, Δ*csrA*, and Δ*fliC*Δ*csrA*. Bars stand for mean ± SEM. * means the significant difference of experimental strain compared to CD1015/E or CD1015 (\**P* < 0.05, \*\**P* < 0.01). One-way ANOVA with post-hoc Tukey test was used for statistical significance.

