## Supplemental Fig. 1-6 and Table S1-3 for "Control of *Clostridiodes difficile* virulence and physiology by the flagellin homeostasis checkpoint FliC-FliW-CsrA in the absence of motility"

**SUPPLEMENTAL MATERIAL**

**Supplementary Figures**

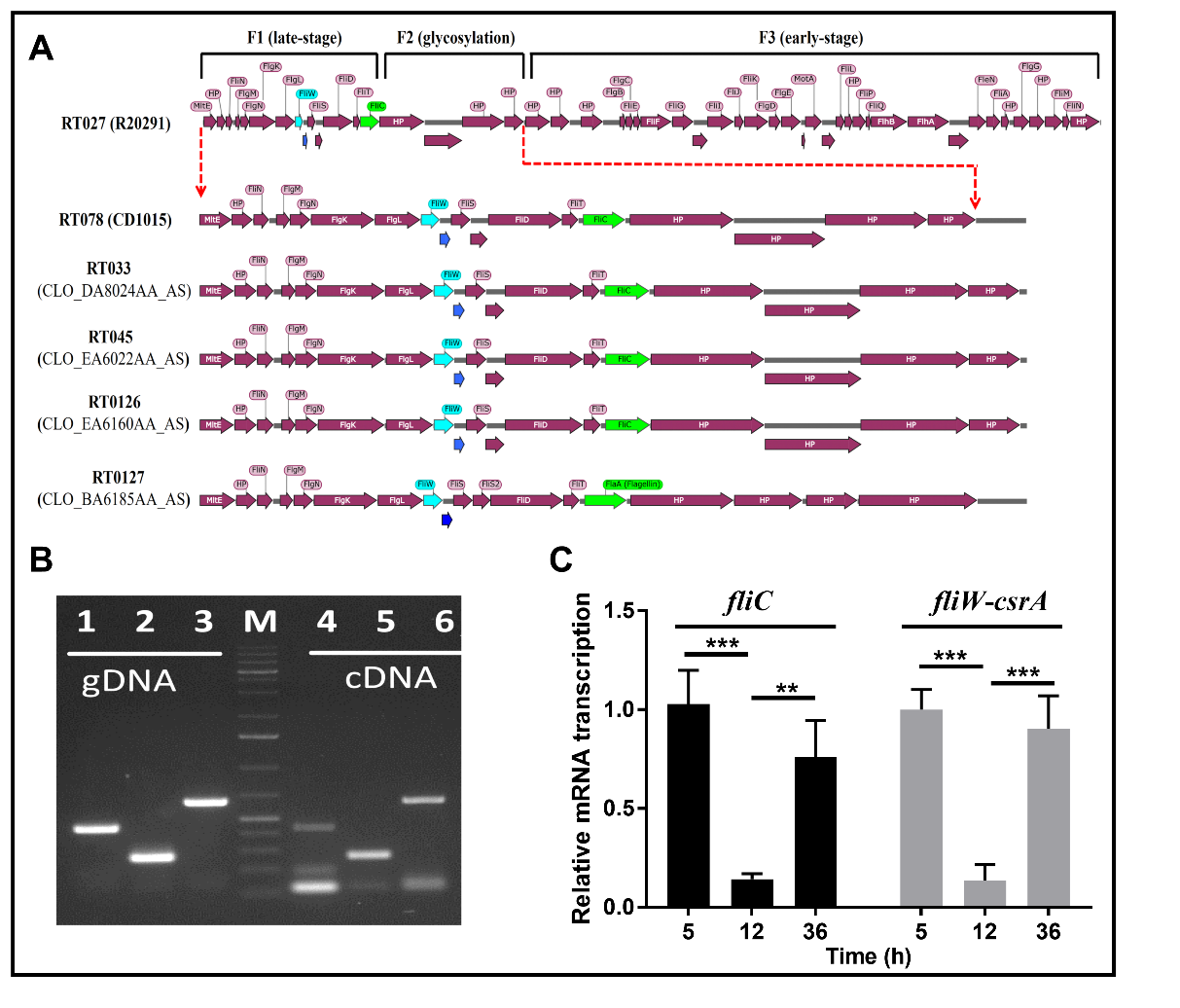

**Fig. S1 Conservation of flagellar genes in non-motile clade 5 strains and verification of *fliW*-*csrA* and *fliC* transcription.**

**(A)** Schematic representation of flagellar genes in the non-motile clade 5 strains and motile RT027 R20291. **(B)** Verification of *fliW* and *csrA* co-transcription by RT-PCR. M: DNA ladder; 1-3: CD1015 genomic DNA was used as PCR template; 4-6: CD1015 cDNA was used as PCR template. 1 and 4: *fliW* PCR test; 2 and 5: *csrA* PCR test; 3 and 6: *fliW*-*csrA* PCR test. **(C)** Transcription analysis of *fliC* and *fliW*-*csrA* in CD1015. Bars stand for mean ± SEM. Differences were considered statistically significant if *P* < 0.05 (***P* < 0.01, ****P* < 0.001). One-way ANOVA with post-hoc Tukey test was used for statistical significance.

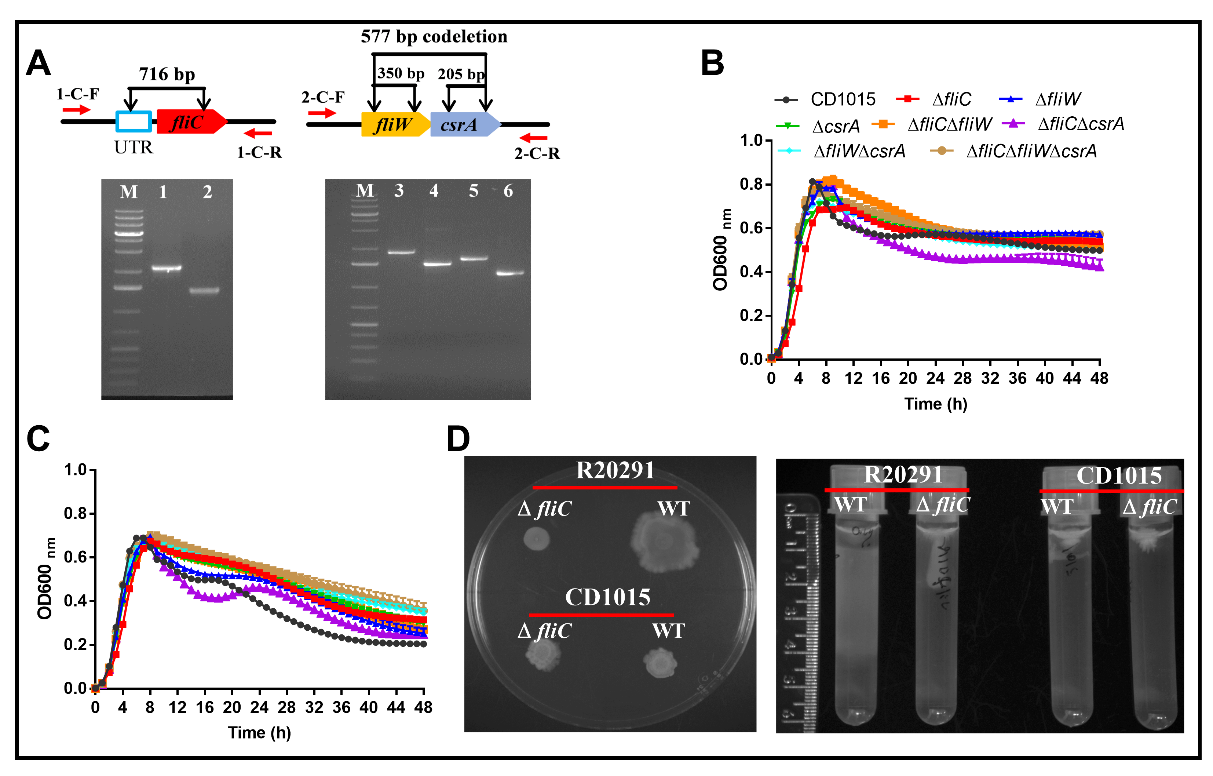

**Fig. S2 Generation of CD1015 derivative mutants, and test of bacterial growth profiles and motility.**

**(A)** Deletion of UTR-*fliC*, *fliW*, *csrA*, and *fliW*-*csrA*. 1-C-F/R were used to verify *fliC* deletion and 2-C-F/R were used to test *fliW*, *csrA*, and *fliW*-*csrA* deletion. M: DNA ladder; 1 and 3: CD1015 genome as PCR template; 2: CD1015∆*fliC* (∆*fliC*) genome test; 4: CD1015∆*fliW* (∆*fliW*) genome test; 5: CD1015∆*csrA* (∆*csrA*) genome test; 6: CD1015∆*fliW*∆*csrA* (∆*fliW*∆*csrA*) genome test. **(B)** Growth profile in BHIS media. **(C)** Growth profile in TY media. **(D)** Bacterial motility test. Swarming and swimming analysis were tested with soft BHIS agar (0.2%) plates and BHIS agar (0.15%) tubes, respectively.

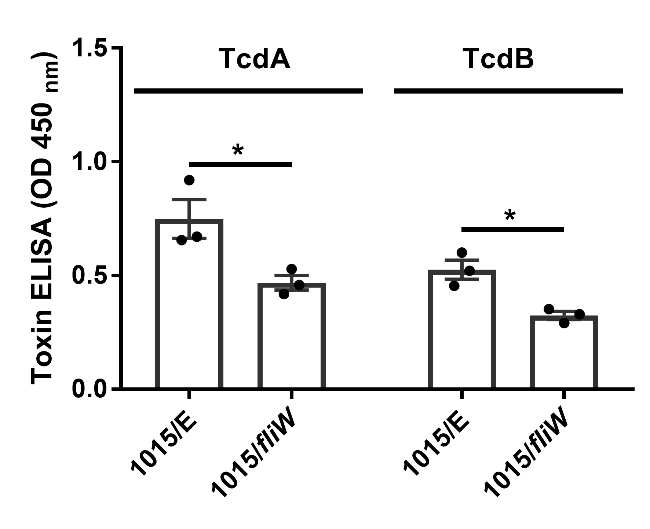

**Fig. S3. Toxin expression in *fliW* overexpression strain.**

TcdA and TcdB concentrations in the supernatants of CD1015-pMTL84153 (1015/E) and *fliW* overexpression strain (1015/*fliW*) were detected by ELISA. Bars stand for mean ± SEM. Differences were considered statistically significant if *P* < 0.05 (**P* < 0.05). Statistical analysis was performed using an unpaired two-tailed *t* test.

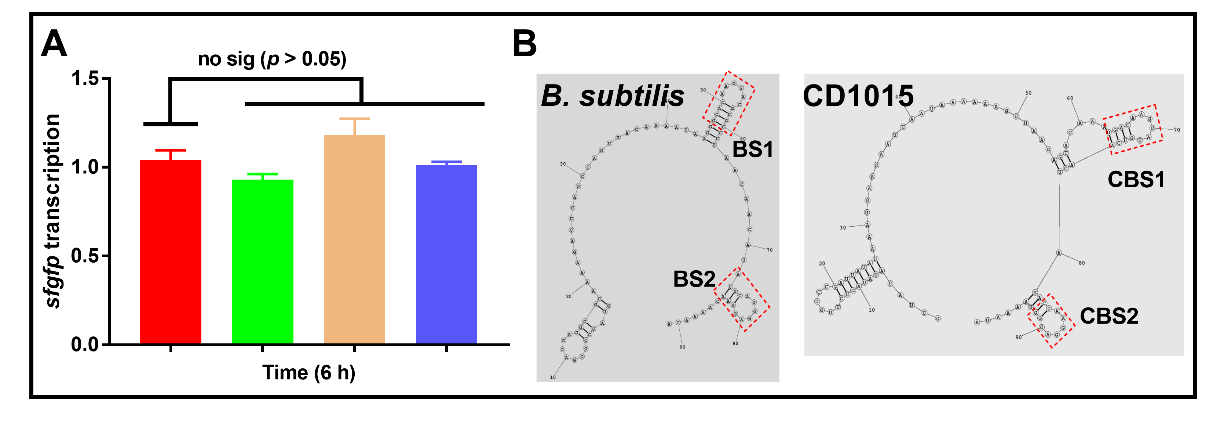

**Fig. S4 Characterization of FliC-FliW-CsrA regulation loop in *C. difficile.***

**(A)** Transcription of *sfgfp* in the different recombinant reporter strains. Red column: P*_lacZ_*-UTR-*sfgfp*; Green column: P*_lacZ_*-UTR-*sfgfp*-P*_tet_*-*fliW*; Orange column: P*_lacZ_*-UTR-*sfgfp*-P*_tet_*-*csrA*; Blue column: P*_lacZ_*-UTR-*sfgfp*-P*_tet_*-*fliW*-*csrA*; **(B)** Structure comparison of *B. subtilis* 5’-UTR of *hag* and CD1015 5’-UTR of *fliC*. BS1 and BS2: CsrA binding sites 1 and 2 in *B. subtilis*. CBS1 and CBS2: CsrA potential binding sites 1 and 2 in CD1015. RNAstructure [dynalign](https://rna.urmc.rochester.edu/RNAstructureWeb/Servers/dynalign/dynalign.html) results were calculated (calculate the lowest free energy secondary structures common to two unaligned sequences).

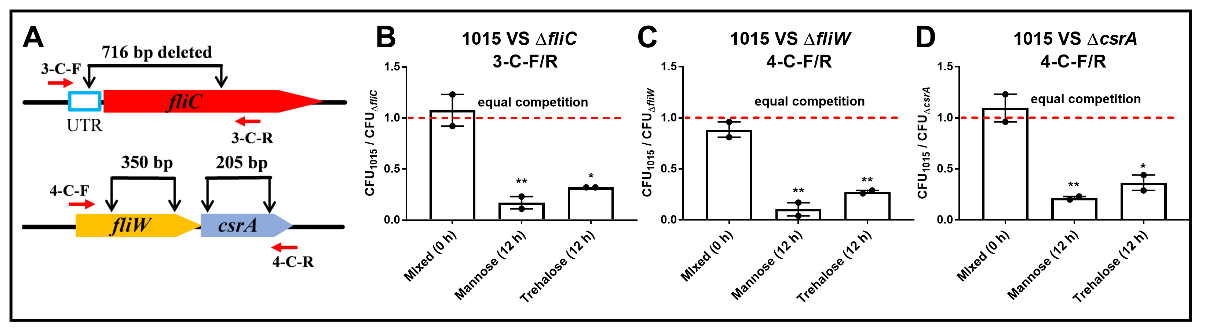

**Fig. S5 ∆*fliC*, ∆*fliW*, and ∆*csrA* mutants outcompete CD1015 in the presence of trehalose and mannose.**

**(A)** Primers used for the colony PCR in the competition test. **(B-D)** CFU_1015_/CFU_mutant_ in the competition cultures. * means the significant difference of CD1015 CFU in the competition mixtures compared to that of at 0 h (**P* < 0.05, ***P* < 0.01). One-way ANOVA with post-hoc Tukey test was used for statistical significance.

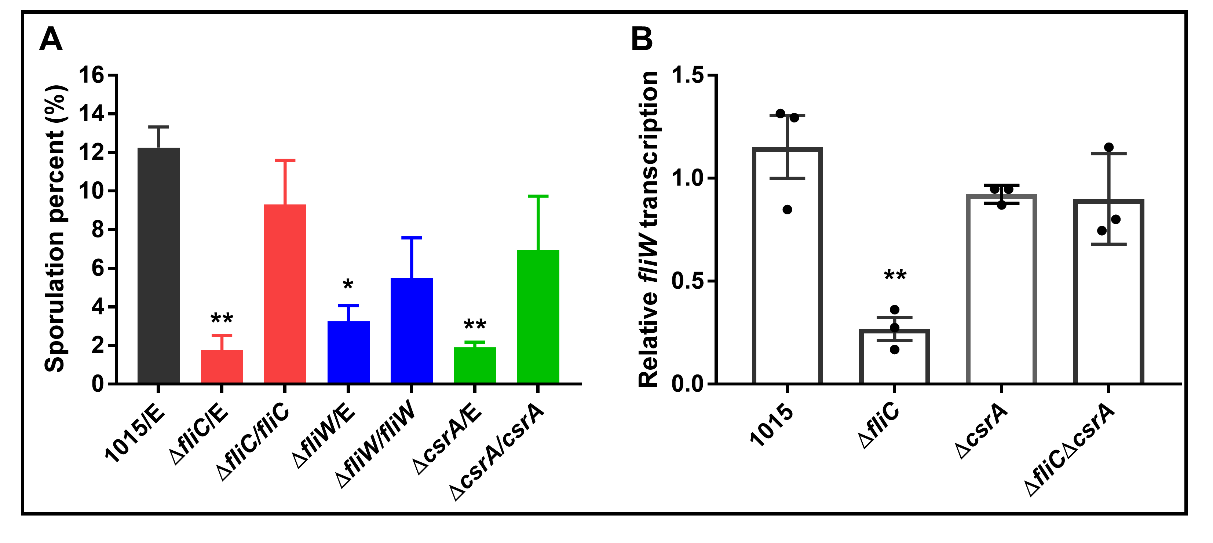

**Fig. S6 Analysis of sporulation and *fliW* transcription in CD1015 and its derivative mutants.**

**(A)** Sporulation analysis. *C. difficile* strains were cultured on 70:30 sporulation agar plates for 5 days, following which the scraped cultures were 10-fold diluted and plated on BHIS plates with 0.1% TA to detect sporulation ratio. The sporulation ratio was calculated as CFU (65 ℃ heated, 30 min) / CFU (not heated). **(B)** Comparison of *fliW* transcription in CD1015, ∆*fliC*, ∆*csrA*, and ∆*fliC*∆*csrA*. Bars stand for mean ± SEM. * means the significant difference of experimental strain compared to CD1015/E or CD1015 (**P* < 0.05, ***P* < 0.01). One-way ANOVA with post-hoc Tukey test was used for statistical significance.

**Supplementary Tables**

**Table S1. Function of late-stage (F1) flagellar genes**

| **Protein** | **Function** | **Reference** |
| --- | --- | --- |
| FliN | part of motor switch | (1) |
| FlgM | inhibitor of FliA regulating late gene expression | (1) |
| FlgN | part of cytoplasmic chaperone | (1) |
| FliS1 | part of cytoplasmic chaperone | (1) |
| FliS2 | part of cytoplasmic chaperone | (1) |
| FliT | part of cytoplasmic chaperone | (1) |
| FlgK | hook-filament Junction | (1) |
| FlgL | hook-filament Junction | (1) |
| FliW | regulating bacterial motility and other physiology phenotypes | (2-4) |
| CsrA | carbon storage regulator | (2-4) |
| FliD | filament cap | (1) |
| FliC | filament protein | (1) |

**Table S2. Bacteria and plasmids utilized in this study**

| **Strains or plasmids** | **Genotype** | **Reference** |
| --- | --- | --- |
| **Strains** |  |  |
| *E. coli* DH5α | Cloning host | NEB |
| *E. coli* BL21 | Protein expression host | NEB |
| *E. coli* BTH101 | Protein-protein interaction analysis host | (5) |
| *E. coli* HB101/pRK24 | Conjugation donor | (6) |
| *C. difficile* 1015 | Clinical isolate; ribotype 078 | (7) |
| *C. difficile* R20291 | Clinical isolate; ribotype 027 | (8) |
| 1015/E | 1015 containing blank plasmid pMTL84153 | This work |
| 1015/*fliW* | 1015 containing pMTL84153-*fliW* | This work |
| R20291Δ*fliC* | R20291 deleted *fliC* gene | This work |
| Δ*fliC* | CD1015 deleted *fliC* gene | This work |
| Δ*fliC/*E | Δ*fliC* containing pMTL84153 | This work |
| Δ*fliC/fliC* | Δ*fliC* containing pMTL84153-*fliC* | This work |
| Δ*fliW* | CD1015 deleted *fliW* gene | This work |
| Δ*fliW/*E | Δ*fliW* containing pMTL84153 | This work |
| Δ*fliW/fliW* | Δ*fliW* containing pMTL84153-*fliW* | This work |
| Δ*csrA* | CD1015 deleted *csrA* gene | This work |
| Δ*csrA/*E | Δ*csrA* containing pMTL84153 | This work |
| Δ*csrA/csrA* | Δ*csrA* containing pMTL84153-*csrA* | This work |
| Δ*fliC*Δ*fliW* | CD1015 deleted *fliC* and *fliW* gene | This work |
| Δ*fliC*Δ*fliW/*E | Δ*fliC*Δ*fliW* containing pMTL84153 | This work |
| Δ*fliC*Δ*fliW/fliW* | Δ*fliC*Δ*fliW* containing pMTL84153-*fliW* | This work |
| Δ*fliC*Δ*csrA* | CD1015 deleted *fliC* and *csrA* gene | This work |
| Δ*fliW*Δ*csrA* | CD1015 deleted *fliW* and *csrA* gene | This work |
| Δ*fliW*Δ*csrA/*E | Δ*fliW*Δ*csrA* containing pMTL84153 | This work |
| Δ*fliW*Δ*csrA/fliW* | Δ*fliW*Δ*csrA* containing pMTL84153-*fliW* | This work |
| Δ*fliC*Δ*fliW*Δ*csrA* | CD1015 deleted *fliC*, *fliW*, and *csrA* gene | This work |
| Δ*fliC*Δ*fliW*Δ*csrA/*E | Δ*fliC*Δ*fliW*Δ*csrA* containing pMTL84153 | This work |
| Δ*fliC*Δ*fliW*Δ*csrA/fliW* | Δ*fliC*Δ*fliW*Δ*csrA* containing pMTL84153-*fliW* | This work |
| **Plasmids** |  |  |
| pKNT25 | T25 reporter at C-terminal | (5) |
| pUT18 | T18 reporter at C-terminal | (5) |
| pKT25 | T25 reporter at N-terminal | (5) |
| pUT18C | T18 reporter at N-terminal | (5) |
| pKNT25-FliW (FliW-T25) | *fliW* fused into N-termnal of T25 reporter | This work |
| pKNT25-FliC (FliW-T25) | *fliC* fused into N-termnal of T25 reporter | This work |
| pKNT25-CrA (CsrA-T25) | *csrA* fused into N-termnal of T25 reporter | This work |
| pUT18-FliW(FliW-T18) | *fliW* fused into N-termnal of T18 reporter | This work |
| pUT18-FliC (FliC-T18) | *fliC* fused into N-termnal of T18 reporter | This work |
| pUT18-CsrA (CsrA-T18) | *csrA* fused into N-termnal of T18 reporter | This work |
| pUT18-TcdR (TcdR-T18) | *tcdR* fused into N-termnal of T18 reporter | This work |
| pET21b | Protein expression plasmid | NEB |
| pET21b-UTR-*sfgfp*-P*_tet_*-*csrA* | Reporter part 5’UTR-*sfgfp* and regulation part P_tet_-*csrA* assembled into pET21b | This work |
| pET21b-UTR-*sfgfp*-P*_tet_*-*fliW* | Reporter part 5’UTR-*sfgfp* and regulation part P*_tet_*-*fliW* assembled into pET21b | This work |
| pET21b-UTR-*sfgfp*-P*_tet_*-*fliW*-*csrA* | Reporter part 5’UTR-*sfgfp* and regulation part P*_tet_*- *fliW*-*csrA* assembled into pET21b | This work |
| pET21b-UTR_CBS1_-*sfgfp*-P*_tet_*-*csrA* | Only kept potential binding site 1 in UTR | This work |
| pET21b-UTR_CBS2_-*sfgfp*-P*_tet_*-*csrA* | Only kept potential binding site 2 in UTR | This work |
| pET21b-UTR_DD_-*sfgfp*-P*_tet_*-*csrA* | Two potential binding sites were deleted | This work |
| pMTL84153 | Complementation plasmid | (9) |
| pMTL84153-*fliC* | pMTL84153 containing *fliC* genes | This work |
| pMTL84153-*fliW* | pMTL84153 containing *fliW* gene | This work |
| pMTL84153-*csrA* | pMTL84153 containing *crsA* gene | This work |
| pDL1 | AsCpfI based gene deletion plasmid | (10) |
| pUC57-PsRNA | sRNA promoter template | This work |
| pDL1-*fliC* | *fliC* gene deletion plasmid | This work |
| pDL1-*fliW* | *fliW* gene deletion plasmid | This work |
| pDL1-*csrA* | *csrA* gene deletion plasmid | This work |
| pDL1-*fliW-csrA* | *fliW-csrA* gene deletion plasmid | This work |

**Table S3. Primers utilized in this study**

| **Primer** | **Sequence (5’ to 3’)** |
| --- | --- |
| 1-F | AGAGATACAGATGTTGCTTCA |
| 1-R | TCCTTGTGGTTGCTGATTA |
| 2-F | AGATGAAGCAGTACTTATAGGA |
| 2-R | GCTTATATTGTTAGGTGCTGATA |
| 3-F | ATGATGAAGGTTACATTAAAAAAAG |
| 3-R | CTAGCATCCACTATCACCTCTC |
| 4-F | ATGCTAGTAATTTCAAGAAAAAAAG |
| 4-R | TTATTTTAATGACTTTAAAATATTT |
| 5-F | GCATGCCTGCAGGTCGACTCTAGAGATGAGAGTTAATACAAATGTAAGTGCT |
| 5-R | AATTCGAGCTCGGTACCCGGGGATCTCCTAATAATTGTAAAACTCCTTGT |
| 6-F | ACGCCACTGCAGGTCGACTCTAGAGATGAGAGTTAATACAAATGTAAGTGCT |
| 6-R | AATTCGAGCTCGGTACCCGGGGATCTTATCCTAATAATTGTAAAACTCCTTGT |
| 7-F | GCATGCCTGCAGGTCGACTCTAGAGATGATGAAGGTTACATTAAAAAAAG |
| 7-R | AATTCGAGCTCGGTACCCGGGGATCGCATCCACTATCACCTCTCAAT |
| 8-F | GCGGGCTGCAGGGTCGACTCTAGAGATGATGAAGGTTACATTAAAAAAAG |
| 8-R | TTAGTTACTTAGGTACCCGGGGATCTTAGCATCCACTATCACCTCTCAAT |
| 9-F | GCATGCCTGCAGGTCGACTCTAGAGATGCTAGTAATTTCAAGAAAAAAA |
| 9-R | AATTCGAGCTCGGTACCCGGGGATCTTTTAATGACTTTAAAATATTTAT |
| 10-F | ACGCCACTGCAGGTCGACTCTAGAGATGCTAGTAATTTCAAGAAAAAAA |
| 10-R | AATTCGAGCTCGGTACCCGGGGATCTTATTTTAATGACTTTAAAATATTTAT |
| 11-F | TTTAACTTTAAGAAGGAGATATACATATGCGTAAAGGCGAAGAGC |
| 11-R | TGTCGACGGAGCTCGAATTCGGATCCTTAGTGGTGATGGTGATGGTGTTTGTACAGTTCATCCATACCA |
| 12-F | CTTTAAGAAGGAGATATACATCTTGTCCGATTATATAAATGAAGAATTAATAAAAAAGTTAAGAGTAGAAATGACAAGGATGTCAACTATACTAAGGAGGGTAAAATA |
| 12-R | CCTTCTTAAAGTTAAACAAATATTTTACCCTCCTTAGTATAGTTGACATCCTTGTCATTTCTACTCTTAACTTTTTTATTAATTCTTCATTTATATAATCGGACAAGA |
| 13-F | CTTTAAGAAGGAGATATACATCTTGTCCGATTATATAAATGAAGAATTAATAAAAAAGTTAAGAGTAGAAA CAACTATACTAAGGAGGGTAAAATA |
| 13-R | CCTTCTTAAAGTTAAACAAATATTTTACCCTCCTTAGTATAGTTGTTTCTACTCTTAACTTTTTTATTAATTCTTCATTTATATAATCGGACAAGA |
| 14-F | CTTTAAGAAGGAGATATACATCTTGTCCGATTATATAAATGAAGAATTAATAAAAAAGTTAAGAGTAGAAATGACAAGGATGTCAACTATATAAAATA |
| 14-R | CCTTCTTAAAGTTAAACAAATATTTTATATAGTTGACATCCTTGTCATTTCTACTCTTAACTTTTTTATTAATTCTTCATTTATATAATCGGACAAGA |
| 15-F | CTTTAAGAAGGAGATATACATCTTGTCCGATTATATAAATGAAGAATTAATAAAAAAGTTAAGAGTAGAAACAACTATATAAAATA |
| 15-R | CCTTCTTAAAGTTAAACAAATATTTTATATAGTTGTTTCTACTCTTAACTTTTTTATTAATTCTTCATTTATATAATCGGACAAGA |
| 16-F | TGGACAGCAAATGGGTCGGGATCCGCATAAAAATAAGAAGCCTGCATTT |
| 16-R | AAAAACCTCCTTTACTGCAGGAGCTCAGATCTGTTAACG |
| 17-F | AGCTCCTGCAGTAAAGGAGGTTTTTATGATGAAGGTTACATTAAAAAAAGG |
| 17-R | CAGTGGTGGTGGTGGTGGTGCTCGATTACTAGCATCCATTATCACC |
| 18-F | AGCTCCTGCAGTAAAGGAGGTTTTTATGCTAGTAATTTCAAGAAAAAAAGATGAAGC |
| 18-R | CAGTGGTGGTGGTGGTGGTGCTCGATTATTTTAATGACTTTAAAATTTTTAT |
| 19-F | AGCTCCTGCAGTAAAGGAGGTTTTTATGATGAAGGTTACATTAAAAAAAGG |
| 19-R | CAGTGGTGGTGGTGGTGGTGCTCGATTATTTTAATGACTTTAAAATTTTTAT |
| 20-F | TGTGTTACATATGACCATGATTACGATGAGAGTTAATACAAATGTAAGTG |
| 20-R | CGCGTGACGTCGACTCTAGAGGATCTTATCCTAATAATTGTAAAACTCCT |
| 21-F | TGTGTTACATATGACCATGATTACGATGATGAAGGTTACATTAAAAAAAG |
| 21-R | CGCGTGACGTCGACTCTAGAGGATCCTAGCATCCACTATCACCTCTC |
| 22-F | TGTGTTACATATGACCATGATTACGATGCTAGTAATTTCAAGAAAAAAAG |
| 22-R | CGCGTGACGTCGACTCTAGAGGATCTTATTTTAATGACTTTAAAATATTT |
| 23-F | AAAGTTAAAAGAAGAAAATAGAAATATAATCTTTAATTTGAAAAGATTTA |
| 24-R | GATAGCTGATGAGTTGTTACAATATCTACAAGAGTAGAAATTAGTATCTAGTTTAGATGTAGCTCTATCTACAAGAGTAGAAATTAATGGT |
| 25-F | ATATTGTAACAACTCATCAGCTATCTAATTTCTACTCTTGTAGATGTATAGAAAAAACTGCTTCTGC |
| 25-R | ATTAATTCTTCATTTATATAATCGG |
| 26-F | CCGATTATATAAATGAAGAATTAATATTAGGGGCACAACAAAATAGA |
| 26-R | CATGCTGATCTAGATTTCTCCATAGATCAATCAGTATTTCGTCTGCAT |
| 1-C-F | TCACTAGCAGGCTATTCTTCG |
| 1-C-R | TAAAAAGTCTAAGCACTGAACAAT |
| 27-R | GATAGCTGATGAGTTGTTACAATATCTACAAGAGTAGAAATTATCCATCAAGTAATTTCTTACCATATCTACAAGAGTAGAAATTAATGGT |
| 28-R | CATGCTGATCTAGATTTCTCCATAGGAATCAATTAGTATTTCATCTGCAT |
| 29-R | TATACCTTCTTCTTCAGTAGAATATCTACAAGAGTAGAAATTACGTCTGACCAAGAGTAATTATATATCTACAAGAGTAGAAATTAATGGT |
| 30-R | GTACTGCTTCATCTTTTTTTCTTATCTACAAGAGTAGAAATTAAAGATGAAGCAGTACTTATAGGAATCTACAAGAGTAGAAATTAATGGT |
| 31-F | ATATTCTACTGAAGAAGAAGGTATATAATTTCTACTCTTGTAGATTGCAAATGCTTTAAATGCAACA |
| 31-R | ATGTAACCTTCATCATATCTGCG |
| 32-F | AACGCAGATATGATGAAGGTTACATTTATTGAGAGGTGATAGTGGAT |
| 32-R | CATGCTGATCTAGATTTCTCCATAGCTATTACAGGCAACACATTATCTAT |
| 33-F | AACGCAGATATGATGAAGGTTACATAAATAATAAGCTTGGAGGAAGT |
| 33-R | CATGCTGATCTAGATTTCTCCATAGTTAAAAGAGCATCTTTGGATA |
| 34-F | ATAAGAAAAAAAGATGAAGCAGTACTAATTTCTACTCTTGTAGATCAGTTAAAGGAGATTCTGTCACAAA |
| 34-R | ATCCACTATCACCTCTCAATAATGG |
| 35-F | CCATTATTGAGAGGTGATAGTGGATAAATAATAAGCTTGGAGGAAGT |
| 35-R | CATGCTGATCTAGATTTCTCCATAGTTAAAAGAGCATCTTTGGATA |
| 2-C-F | CTTAGTAATGAAACCAATGAAAAAA |
| 2-C-R | CTGCATCTTGGTCTAATGTAGTCAT |
| 36-F | GCAAGTTGAGCGATTTACTTCGGT |
| 36-R | GTACTGGCTCACCTTTGATATT |
| 37-F | GCGGAAATGGTAGAAATG |
| 37-R | ATCAGGTGCTATCAATACTT |
| 38-F | CTTACACCATCTATGATAGTTG |
| 38-R | AAGTTCAAGTTTATGCTCAAT |
| 39-F | CCAAGTGGTACTATACATGCTAT |
| 39-R | TGAGGAACATTCGTAACATCTAT |
| 40-F | AAGAGATAACGGATATGATATAGA |
| 40-R | GTGTTCAGTAGATGTATGGT |
| 41-F | ACCAACCACGCATAATATCTA |
| 41-R | AGTCATACCTATTTCTTCACCTT |
| 42-F | GCTGTCTTCTGGTGTTAG |
| 42-R | TTCTTCCTGCTTGGTCTA |
| 43-F | CTGAAGAAGAAGGTATAGGAT |
| 43-R | CGTCTGACCAAGAGTAAT |
| 3-C-F | AGCTTACTAAACAAGTGAACACAAT |
| 3-C-R | TTATTCCAATTATGAAAATACACTT |
| 4-C-F | GTTCAATTAAAGGCAGCAGAGT |
| 4-C-R | TCCTGAACTCTTACAAGCTCTCTAT |
